# Metastable phase-separated droplet generation and long-time DNA enrichment by laser-induced Soret effect

**DOI:** 10.1101/2024.10.23.619780

**Authors:** Mika Kobayashi, Yoshihiro Minagawa, Hiroyuki Noji

**Affiliations:** Department of Applied Chemistry, Graduate School of Engineering University of Tokyo, Tokyo 113-8656, Japan

**Keywords:** LLPS, laser-induced phase separation, droplet kinetics, DNA enrichment, thermophoresis

## Abstract

Spatiotemporally controlled laser-induced phase separation (LIPS) offers unique research avenues and has a potential for biological and biomedical applications. However, LIPS conditions often have drawbacks for practical use, which limit their applications. For instance, LIPS droplets are tentative and diminish after the laser is terminated. Here, we developed a novel LIPS method using laser-induced Soret effect with a simple setup to solve these problems. We generate liquid-liquid phase-separated (LLPS) droplets using LIPS in an aqueous two-phase system (ATPS) of dextran (DEX) and polyethylene glycol (PEG). When DEX-rich droplets were generated in the DEX/PEG mix on the phase boundary, the droplets showed unprecedently high longevity; the DEX droplets were retained over 48 h. This counterintuitive behavior suggests that the droplet is in an unknown metastable state. By exploiting the capability of DEX-rich droplets to enrich nucleic acid polymers, we achieved stable DNA enrichment in LIPS DEX droplets with a high enrichment factor of 1400 ± 400. Further, we patterned DNA-carrying DEX-rich droplets into a designed structure to demonstrate the stability and spatiotemporal controllability of DEX-rich droplet formation. This is the first report for LIPS droplet generation in a DEX/PEG system with flexible conditions for the usage of LIPS droplets, opening new avenues for biological and medical applications of LIPS.

---

Liquid-liquid phase separation (LLPS) attracts large interests in biological sciences since the recent findings that phase separation plays important roles in a wide variety of biomolecular systems(1–4). In addition to biological droplets, the ATPS (aqueous two-phase system) of Dextran (DEX) and polyethylene glycol (PEG) also draws a large attention because ATPS of DEX/PEG can enrich some of biomolecules in DEX-rich phase(5–8) while these polymers are chemically and biologically inert and thereby they show fine biocompatibility. In particular, DEX-rich droplets in DEX/PEG ATPS have been reported to well enrich DNA molecules(9). Other biomolecules such as RNA and proteins are also reported to have enriched in the DEX-rich phase(10, 11). For these reasons, DEX/PEG ATPS was utilized in microchemical/biological systems for efficient DNA and protein enrichment(8, 12) or in artificial cell models to enrich biomolecules and/or enhance biomolecular reactions(11, 13–15). Thus, spatiotemporal control of LLPS including DEX/PEG ATPS under the microscope is of great importance for biological applications.

Laser-induced phase separation (LIPS) is a method to generate LLPS droplets in a controlled manner and it has been investigated in several systems(16–27). Local heating upon laser irradiation to LCST (lower critical solution temperature) systems is a simple LIPS method(19) where droplets are created when the heated region satisfies the two-phase condition. However, the generation temperature for the LIPS in LCST systems is limited by the phase diagram and generally it requires high generation temperature which is not applicable in biological experiments. Especially, it is difficult to keep precise temperature control for biological reactions. In the case of DEX/PEG systems, there are some systems which show LCST(28). In principle, LIPS in those systems should be possible but LIPS DEX droplet generation has never been reported so far. This would be partly because of the above reasons.

Another approach for LIPS is to use the Soret effect (thermophoresis), which is a non-equilibrium phenomenon wherein a concentration gradient is formed under a temperature gradient(29). In this case, local concentration is modulated by temperature gradient, but temperature does not directly drive phase separation. This means that the generation temperature is basically not limited by the phase diagram. LIPS by the Soret effect can be conducted at a desired temperature if the appropriate concentration gradient is formed. The concentration gradient of a molecule induced by the Soret effect, *∇c*, is described as

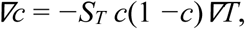

where *c, ∇T*, and *S_T_* represent the molecular fraction, temperature gradient, and ratio of the thermal mass diffusion coefficient (*D_T_*) to the mutual mass diffusion coefficient (*D*), respectively; *S_T_* = *D_T_/D*. The value of *S_T_* ranges from 10^−5^ to 10^−1^ (30). Whether a molecule accumulates (*S_T_ <* 0) or is excluded (*S_T_ >* 0) along a temperature gradient depends on its species and its combination with other molecules in the mixture(31). On the other hand, the concentration gradient generated by the Soret effect is generally not sufficient to induce phase separation. There are, therefore, only a few reports on LIPS using the Soret effect. Delville et al. demonstrated LIPS in a quaternary micellar system because the required concentration change was sufficiently small to achieve phase separation(16, 17). Voit et al. studied thermal patterning on a polymer blend using a focused laser beam and modulated phase separation near the critical point where the Soret coefficient diverges(21, 22).

Another drawback of LIPS so far is that LIPS droplets prepared using these methods are ephemeral and disappear after the termination of laser irradiation because the droplets are formed under non-equilibrium conditions in a stable single phase. The duration of the LIPS droplets is basically determined by the diffusion of molecules. In previous reports on LIPS experiments with binary mixtures, LIPS droplets disappeared within tens of minutes(21, 25). Thus, LIPS droplets are principally unstable and revert to the miscible solution state after laser induction(21, 25), which limits its range of application.

In this study, we aimed to develop a novel LIPS methodology for the generation of phase-separated DEX-rich droplets in a DEX/PEG system by use of the laser-induced Soret effect. DEX has a unique feature; this polymer exhibits a negative Soret coefficient, *S_T_ <* 0(31); DEX molecules accumulate to the hot region along the temperature gradient. Thus, the negative *S_T_* of DEX is, in principle, suitable for LIPS with the Soret effect. We developed a local heating system in which an infrared laser spot was introduced on a coverslip coated with a semiconductor which absorbs the infrared light and produces heat. The local heating produced a temperature gradient, which resulted in the local condensation of DEX via the Soret effect. The laser-induced Soret effect successfully triggered the formation of DEX-rich droplets. We discovered that the generated droplets were extraordinarily stable; the droplets were retained for over 48 h. It was also found that the DEX-rich droplets enriched DNA molecules by a factor of 1400. Finally, we demonstrated the patterning of DNA-containing droplets along the defined structures, by exploiting the controllability, longevity and capability for DNA enrichment of the LIPS-induced DEX-rich droplet.

## LIPS droplet generation by laser-induced Soret effect

We produced LIPS droplets in a miscible mixture of DEX (*Mw* = 4.5– 6.5 × 10^5^) and PEG (*Mw* = 3.5 × 10^4^). A phase diagram of the DEX/PEG system is illustrated in Fig. 1a. It does not change within the experimental temperature range of 293–363 K. Therefore, heating by laser irradiation did not directly affect the phase of the DEX/PEG system (See Methods). We performed experiments on a DEX/PEG mix on the coexisting curve, which is the phase boundary between single- and two-phase states, with the expectation that the system would inevitably enter the two-phase region by the small change in concentration induced by the Soret effect. The DEX/PEG mixture on the coexisting curve was prepared experimentally as follows; at first, a DEX/PEG was prepared at an initial composition in the two-phase region —DEX at 6 wt.% and PEG at 2.6 wt.% (open circles in Fig. 1a). The DEX/PEG mixture was equilibrated to form macroscopic two phases and complete phase separation, as shown in the photograph in Fig. 1a. Each phase obtained by the preparation should be exactly on the phase boundary because it is an equilibrium phase. The DEX/PEG compositions of the upper PEG-rich phase and the lower DEX-rich phase are estimated from the fluorescent intensity of the spiked, fluorescently labelled DEX and PEG (Fig. 1a), respectively. The compositions were consistent with those reported in a previous study(32). The upper PEG-rich phase solution was used in subsequent experiments to induce DEX-rich droplet generation by LIPS.

**Fig. 1:**
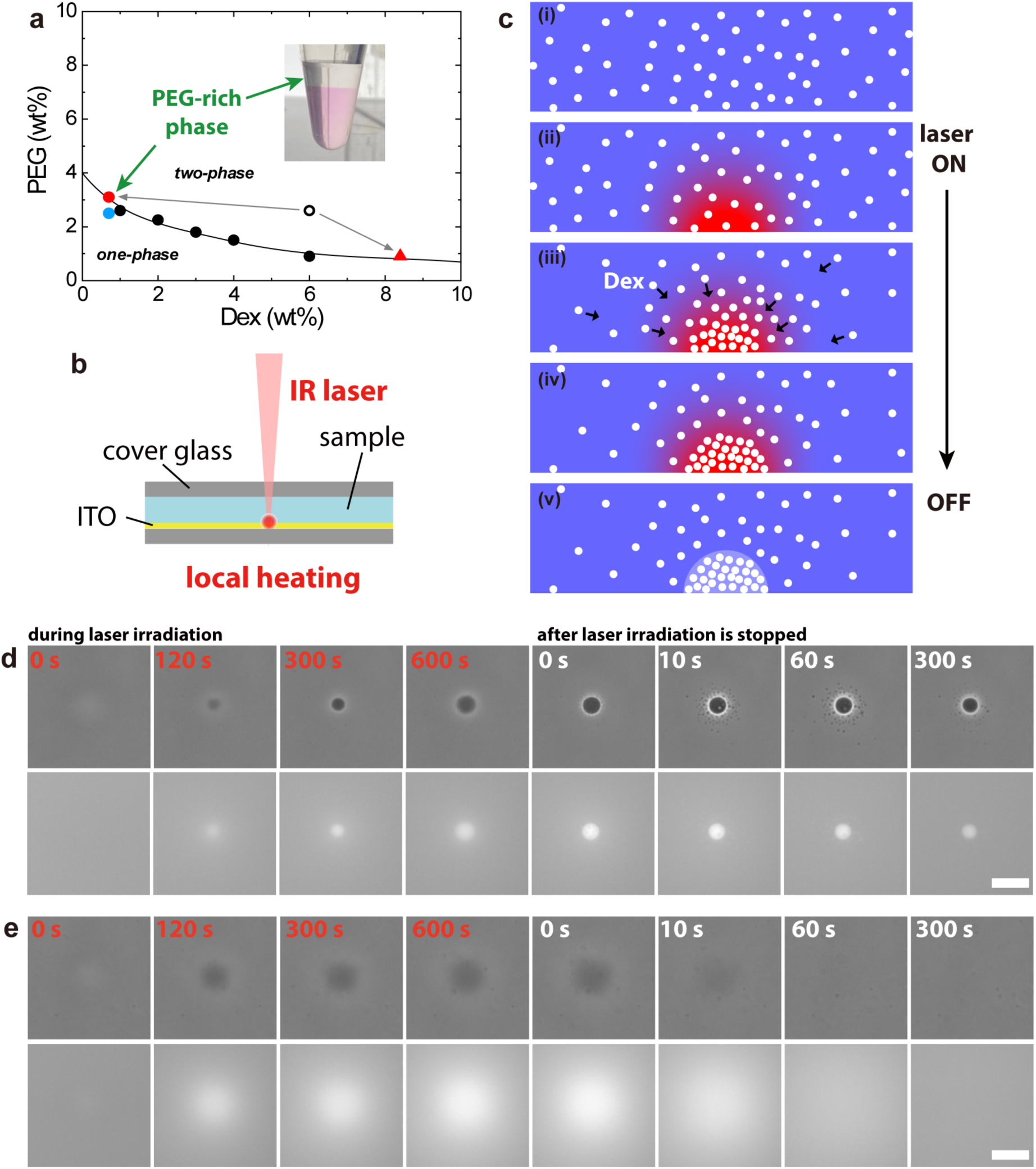
Droplet generation induced by the Soret effect. **a**, Phase diagram of the ATPS of Dextran (*M_w_* = 5.5 × 10^5^) and PEG (*M_w_* = 3.5 × 10^4^). The black circles are phase boundary obtained by visual observation. A sample was prepared with an initial composition of 6 wt.% and PEG 2.6wt.% (open circle) in a two-phase region. The mixture was phase-separated, and the composition of each phase is indicated by a red circle and triangle (See Methods). The upper PEG-rich phase of the phase-separated mixture was used in droplet generation experiments. The black curve represents the coexisting curve. **b**, Experimental setup. The ITO coated on the cover glass absorbs laser light and causes local heating. **c**) Droplet generation process induced by the Soret effect. (See text.) Side view of the laser focus region. The white particles denote the dextran molecules. (i) Before laser irradiation. (ii) A temperature gradient occurs at the ITO surface when the laser irradiation is initiated. (iii) Dextran molecules enter the hot region. (iv) A dextran-rich domain formed. (v) The phase-separated droplets remain after the laser is switched off. **d**, Droplet generation process (Sample: upper phase of DEX 6 wt.% and PEG 2.6 wt.% including fluorescently labelled DEX 0.2 wt.%). (top) Phase contrast images. (Bottom) Fluorescence images of dextran. When the laser irradiation was initiated, a dark dextran-rich domain gradually appeared, and the droplet interface became sharp when the laser irradiation was stopped (see text). Successful droplet generation was observed only for the sample at the phase boundary. Scale bar = 20 μm. **e**, Cases in which no droplets were generated (DEX 0.7 wt.% and PEG 2.5 wt.% blue circle in the phase diagram). The sample composition was very close to that of **d** but only the behavior caused by the Soret effect was observed. Dextran collected by the Soret effect diffuses rapidly after the laser is switched off (local heating is removed). (top) Phase contrast images. (Bottom) Fluorescence images of dextran. Scale bar = 20 μm.

Local heating was performed under a microscope using an infrared (IR) laser. Figure 1b illustrates a schematic of the experimental setup. A temperature gradient was generated at the laser focus point, where the IR laser light was absorbed by the ITO coat on a coverslip of a sample cell. The schematic of DEX-rich droplet generation by LIPS is shown in Fig. 1c. The temperature gradient produced upon the laser irradiation attracts DEX molecules via the Soret effect. The temperature at the heating spot during laser irradiation was estimated to be +10 K from the surrounding medium, generating a temperature gradient of 0.1 K/μm (*SI Appendix*, section 1). Subsequently, a DEX-rich droplet phase appeared at the focal point of the IR laser.

Figure 1d illustrates time-lapsed images of the droplet-generation process (See also *SI Appendix*, Movie 1). When the laser irradiation was initiated, the growth of the dark domain in the phase contrast images was observed, as well as the growth of a fluorescence spot in the fluorescence images. After the irradiation time for 10 min, the laser was switched off. Then, the small droplets suddenly appeared around the main droplet (see phase-contrast images at 10 or 60 s after laser termination in Fig. 1d). These satellite droplets gradually coalesced into the main droplet. The droplet interface was not clear during laser irradiation, and it became sharp after laser termination. It is likely that the active mass transport of DEX by the laser-induced Soret effect perturbed the formation of a stable interface. We also examined LIPS using an ITO-free glass substrate. DEX accumulation and droplet formation were not observed (*SI Appendix*, section 2), confirming that droplet generation was not induced by the radiation pressure or electromagnetic force of the laser, but by local heating, which caused the Soret effect. We confirmed that the temperature of the generated droplet can be changed after the laser is switched off (*SI Appendix*, section 3).

When the DEX/PEG mix was prepared at concentrations below the coexisting curve (blue circle in the phase diagram in Fig. 1a), laser irradiation failed to form a droplet, although DEX condensed (Fig. 1e). When the irradiation was terminated, the accumulated DEX molecules diffused within minutes. This means we need an initial sample very close to the coexisting curve for LIPS using the Soret effect as more investigated below.

## Longevity of LIPS droplet

The generated DEX droplets exhibited remarkable longevity. Figure 2a shows a representative series of time lapsed images for 2 days. Figure 2b exhibits the time course of the diameter of the droplet shown in Fig. 2a. Initially, the diameter was approximately 7 μm. After the initial shrinkage process of 1–2 h, the droplet size reached a steady value of approximately 3 μm. Figure 2c illustrates the diameters of the droplets generated under the same experimental condition plotted as a function of the laser irradiation time *t*_IR_. The initial diameter ranged from 6–12 µm, and the final diameter was 1–3 µm. Neither the initial nor the final diameter did not show a clear *t*_IR_-dependence. This suggests that DEX condensation by the laser-induced Soret effect plateaued within 5 min of irradiation.

**Fig. 2:**
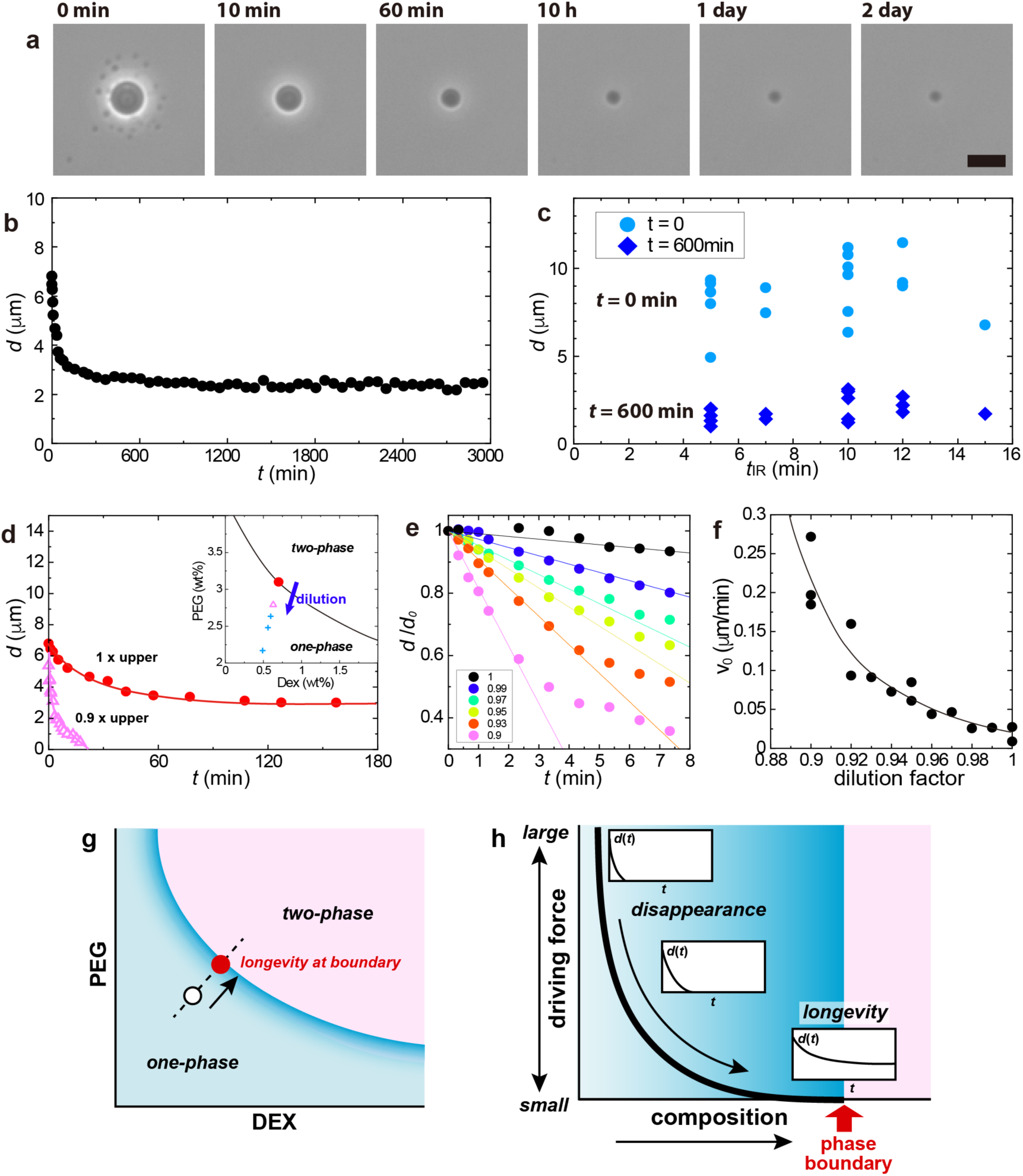
Long-time droplet presence and its relationship with the distance to the coexisting curve. **a-c**, Cases where the droplet generated in the sample on the coexisting curve sustains for a long time. The droplet was generated in the upper phase of 6 wt.% and PEG 2.6 wt.% (red circle in Fig. 1a). (**a**) Time evolution of phase-contrast images after the laser was switched off. (Laser irradiation time 10 min.) The number denotes the time elapsed after the laser was disabled. The droplets did not disappear and remained for more than two days. Scale bar = 10 μm. (**b**) Time evolution of droplet diameter. The value approached a finite value of approximately 3 μm and converged. (**c**); Droplet diameters at *t* = 0 and 600 min plotted against the laser irradiation time *t*_IR_. **d-f**, Relationship between droplet generation behavior and distance to the coexisting curve obtained by dilution experiments on the upper PEG-rich phase (see text). (**d**); Time evolution of droplet diameter when the generated droplet disappears (pink triangles). This was attributed to the laser irradiation experiment using a sample of the diluted upper PEG-rich phase with a dilution factor of 0.9. The long-time presence achieved without dilution (dilution factor 1) is plotted for comparison (red circles). The solid curves are visual guides. The inset shows the schematic illustration of dilution experiments. Red circle: upper PEG-rich phase: sample on the coexisting curve (solid curve). Diluting the upper PEG-rich phase increases the distance from the coexisting curve (a smaller dilution factor corresponds to being more distant from the coexisting curve). The condition where the produced droplet disappeared (Pink triangle: dilution factor = 0.9). Further dilution makes a condition where droplet generation is unsuccessful (Blue crosses). (**e**); Droplet diameter in the initial regime. The diameter is normalized by the initial value at *t* = 0. The lines represent the linear fits of the initial linear regime of time dependence. Numbers in the legend denote the dilution factor. (**f**); Initial speed of droplet shrinkage obtained as the slope in the fit results in **e**. The speed decreases approaching the coexisting curve: dilution factor =1. The solid curves represent visual guides. **g,h**, Schematic illustration of the mechanism of longevity. (**g**); Schematic phase diagram of a DEX/PEG system. LIPS droplets generated from the open circle inside the one-phase region disappear. When the initial sample is on the phase boundary (red circle), LIPS droplet exhibits longevity. (**h**); The relationship between the behavior of a LIPS droplet (time dependence of droplet diameter d(t) after switching off the laser) and the initial composition of the sample. The longevity appears when the initial composition is approaching the phase boundary because of decreasing the driving force to make the created LIPS droplet disappear.

We investigated the relationship between the droplet longevity and the initial composition of DEX/PEG mixture. In particular, we asked how stable the generated droplets are when the initial concentrations of DEX and PEG are below the coexisting curve (inset of Fig. 2d). For this purpose, we prepared DEX/PEG mixtures by diluting the PEG-rich phase taken from DEX/PEG ATPS solution, which systematically changed the distance from the coexisting curve (inset of Fig 2d). When the DEX/PEG mix was prepared at a dilution factor of 0.9 (pink triangle in the inset of Fig. 2d), LIPS droplets were once generated, however, they disappeared within 30 min (pink triangles in Fig. 2d). The droplets prepared in diluted mix shrank significantly faster than that without dilution. When the DEX/PEG mix was more diluted; the dilution factor was less than 0.9 (blue crosses in the inset of Fig. 2d), and no droplet formation was observed. Figure 2e shows the time-courses of the droplet diameter for the samples with the dilution factors from 1 and 0.9. We analyzed the initial linear regime using linear fitting to estimate the initial speed of droplet shrinkage, *v*_0_. The results are shown in Fig. 2f. The rate of shrinkage, *v*_0_ decreases systematically as the dilution factor approaches 1 (no dilution: on the coexisting curve). This suggests that the driving force to revert the droplet to the original miscible state decreases, when the composition of DEX/PEG mix approaches the coexisting curve. The relationship between the initial composition in the phase diagram and the appearance of longevity is schematically explained in Fig. 2 g and h. These findings confirm that the DEX/PEG mix composition has to be quite close or on the coexisting curve for stable DEX droplet generation.

## DNA enrichment in LIPS droplet

We attempted DNA enrichment in LIPS DEX droplets by exploiting the capability of DEX-rich droplets to enrich DNA molecules(9). The concept of our experiment is schematically illustrated in Fig. 3a; the longevity of DEX-rich LIPS droplets is expected to achieve stable DNA enrichment. First, we examined the possibility of DNA enrichment in LIPS droplets. The PEG-rich phase solution for LIPS was prepared with *λ*-DNA (48,502 bps) through the protocol as same as the above experiments. The DNA concentration in the prepared PEG-rich solution was 70 pg/μL (see Methods and *SI Appendix*, section 4). We conducted LIPS with DNA-containing PEG-rich solution as same as above. Fig. 3b shows the images of a LIPS DEX droplet generated upon laser irradiation for 5 min. The fluorescence image of the DNA intercalator reveals the enrichment of DNA in the LIPS droplet. The droplets with enriched DNA were as stable as droplets prepared without DNA. After initial shrinkage, the droplets retained a constant diameter over 24 h. The fluorescence signal from DNA was more stable than that from DEX; the fluorescence signal of DNA in droplets was observed even after 18 h (Fig. 3c), while the fluorescence signal of DEX became significantly darker compared with the initial fluorescence signal (*t* = 0 h).

**Fig. 3:**
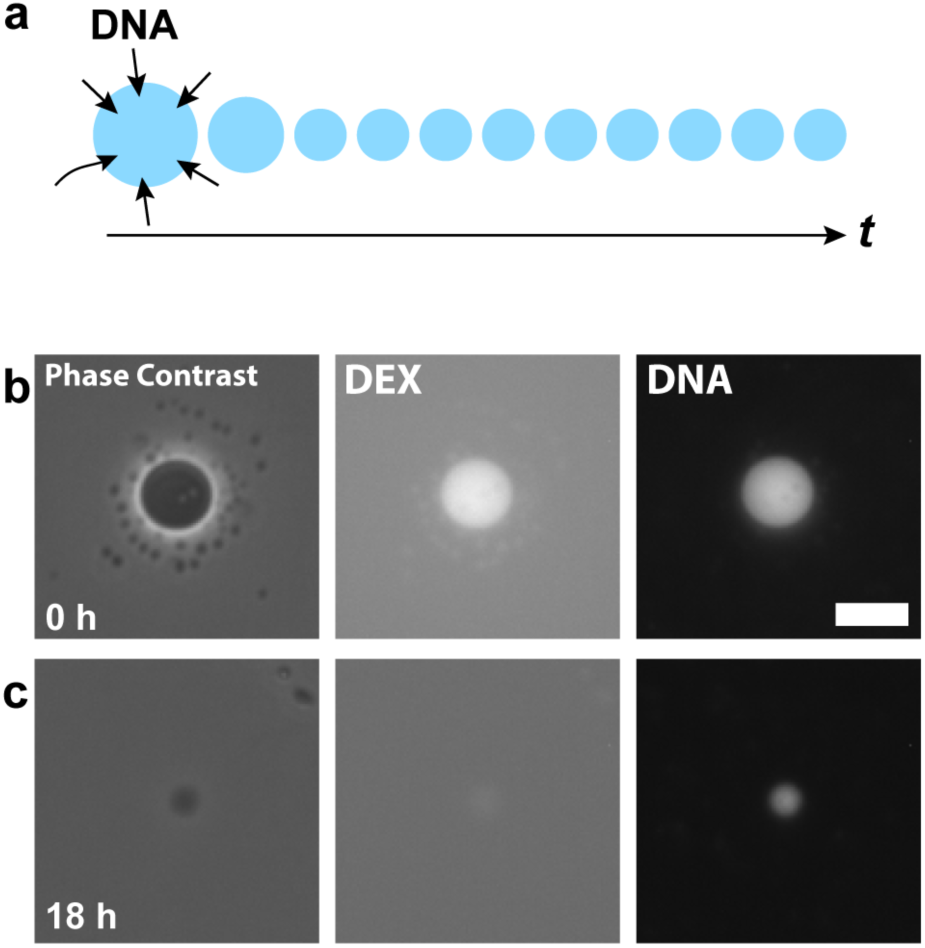
DNA enrichment in the LIPS droplet. **a**, Schematic illustration of the concept of long-time DNA enrichment in LIPS droplets. **b, c**, The droplet was induced by laser irradiation for 5 min to the upper PEG-rich phase of DEX 6 wt.% and PEG 2.6wt.% + *λ*-DNA (48,502 bp) 29 ng/µL. **b**, *t* = 0 h; **c**, *t* = 18 h after the laser is disabled. Phase-contrast image, fluorescent image of dextran, and fluorescent image of the DNA observed by epifluorescence microscopy from left to right. The droplet was visualized using a DNA intercalator and fluorescently labelled dextran. The DNA was successfully incorporated into the generated dextran-rich droplet. Scale bar = 10 μm.

For more quantitative analysis on the concentration of DNA and DEX of the LIPS droplets, the fluorescence intensity of the droplets was measured with confocal microscopy. This is because the fluorescence images of the droplets in epifluorescence microscopy contained a considerable level of background fluorescence intensity from the surrounding medium, hampering accurate measurement. LIPS droplets were prepared on laser-equipped epifluorescence microscopy and subsequently translated to confocal microscopy for time-evolution observation. Therefore, the confocal microscopy observations were started several minutes after LIPS. However, as shown below, the time evolution of the fluorescent signal of DNA/DEX in LIPS droplets exhibited significantly slower change, in the order of tens of minutes. Thus, the delay caused by sample translation was neglected.

Figure 4a displays the representative time-lapsed images of the LIPS droplets visualized by fluorescent DNA/DEX after laser irradiation for 12 min. The time course of droplet diameter is shown in *SI Appendix*, Fig. S1a. It showed an exponential decay and converged to a finite value around 3 μm. The fluorescent intensity of DNA in the LIPS droplet showed a temporal rise around 51 min, followed by decay. In contrast, the fluorescence intensity of DEX showed a simple monotonous decay. We defined the enrichment factor of DNA, *EF*^DNA^ as the ratio of DNA concentration in a LIPS droplet to that in the surrounding PEG-rich solution, *EF*^DNA^ = [DNA]_droplet_/[DNA]_surrounding_. The DNA concentration in droplets ranged from 50 to 200 ng/μL, while that of the surrounding PEG-rich phase solution was 70 ± 10 pg/μL (see Methods and *SI Appendix*, section 4). The time evolution of *EF*^DNA^ is illustrated in Fig. 4b. As observed in the time-lapsed images, *EF*^DNA^ initially showed a transient rise to the maximum value around 50 min, followed by slow decay to a steady level. The statistical analysis showed the mean values of the initial, maximum, and steady stages were 1200, 2000, 1400 (Fig. 4d). The individual data were plotted against the initial droplet diameter *d*_0_ in *SI Appendix*, Fig. S2a. Although the exact mechanism for the transient rise of *EF*^DNA^ is not clear, it is attributable to condensation upon the droplet shrinkage (See *SI Appendix*, Fig. S1a). Time *t*_max_ when *EF*^DNA^ showed its maximum is shown in *SI Appendix*, Fig. S3a. In some cases, there was no evident maxima (*SI Appendix*, Fig. S3b).

**Fig. 4:**
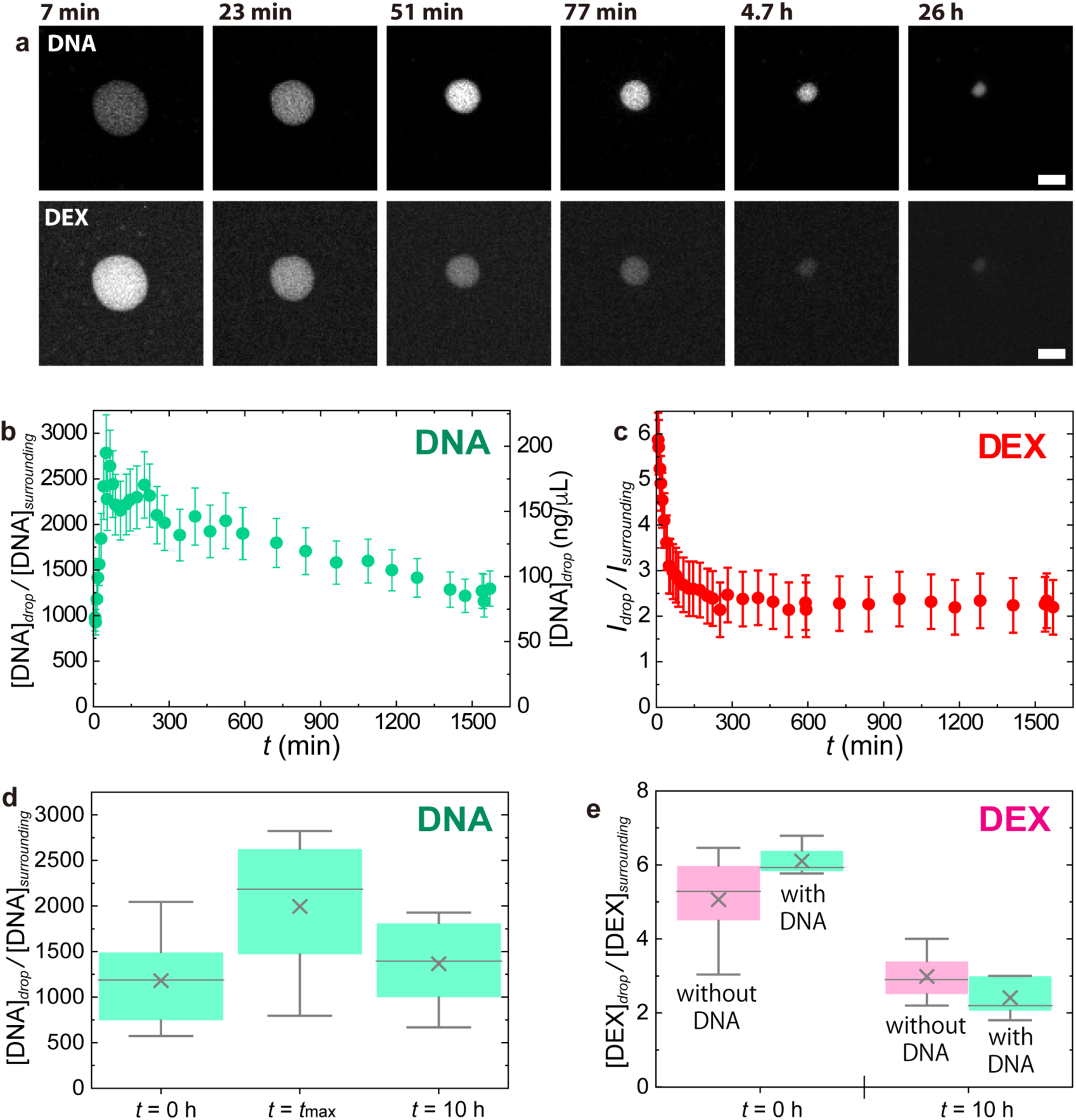
Time evolution of LIPS droplet observed by confocal measurements. The droplet was generated by laser irradiation for 12 min to the upper PEG-rich phase of DEX 6 wt.% and PEG 2.6 wt.% + *λ*-DNA (48,502 bp) 29 ng/µL. **a**, Time-lapse image of a droplet visualized using a DNA intercalator and fluorescently labelled dextran. The time in each image is the time after the termination of the laser. Scale bar = 5 μm. **b**, DNA enrichment factor. Error bars are estimated from standard deviation in the estimation of *c*_surrounding_. **c**, Fluorescence intensity ratio of dextran. Error bars represent uncertainties from focal positions. **d**, DNA enrichment factor at characteristic times. The time *t*_max_ denotes the time when the DNA enrichment factor reaches its maximum value. It depends on the experiment (51 min for Fig. 4b) and is shown in *SI Appendix*, Fig. S3. **e**, Dex intensity ratio at characteristic times in the case with and without DNA. In **d** and **e**, 25–75 percentile (box), median (line), mean (cross), highest/lowest observations (whiskers).

The concentration ratio of DEX of the droplet to the surrounding solution, *CR*^DEX^ was estimated as the fluorescence intensity ratio of the droplet to that in the surrounding PEG-rich solution, *CR*^DEX^ = [DEX]_droplet_/[DEX]_surrounding_ (*SI Appendix*, section 4). Fig. 4c shows the time evolution of *CR*^DEX^that exhibited an exponential decay from approximately 6 to a plateau level, around 2 (Fig. 4c). The trend of the DEX concentration decrease can be seen in the result of statistical analysis shown in Fig. 4e. The validity of the change is discussed in *SI Appendix*, section 5. The individual data were plotted against the initial droplet diameter *d*_0_ in *SI Appendix*, Fig. S2b. The timescale of the decay in *CR*^DEX^ was similar to that of the diameter (*SI Appendix*, Fig. S4).

The presence of DNA did not show a significant effect on the behavior of the LIPS droplets. The decay rate of the droplet diameter was the order of 10^1^ min in both cases, w/ and w/o DNA (*SI Appendix*, Fig. S1b). The initial value and the final value of *CR*^DEX^ were almost the same between w/ and w/o DNA shown in Fig. 4e.

## Comparison of LIPS droplet and spontaneous droplet

We compared LIPS droplets with droplets formed by spontaneous phase separation. The spontaneous droplets of the DEX-rich phase were prepared by agitation of the two-phase DEX/PEG mix which is the same composition as that used in LIPS experiments and observed in confocal microscopy (*SI Appendix*, section 6). The DNA enrichment factor *EF*^DNA^ and the DEX concentration ratio *CR*^DEX^ of spontaneous droplets were compared with those of LIPS droplets in Figs. 5a and 5b, respectively. The mean values of initial or final *EF*^DNA^ for LIPS droplets are 1200 and 1400 which are higher than *EF*^DNA^ for spontaneous droplets, around 600 (Fig.5a). The distribution of *EF*^DNA^ for LIPS droplets is wider than that of spontaneous ones. The mean values of initial or final *CR*^DEX^ for LIPS droplets are 6.9 and 3.0 which are smaller than *CR*^DEX^ of spontaneous ones, 9.4 (Fig.5b).

**Fig. 5:**
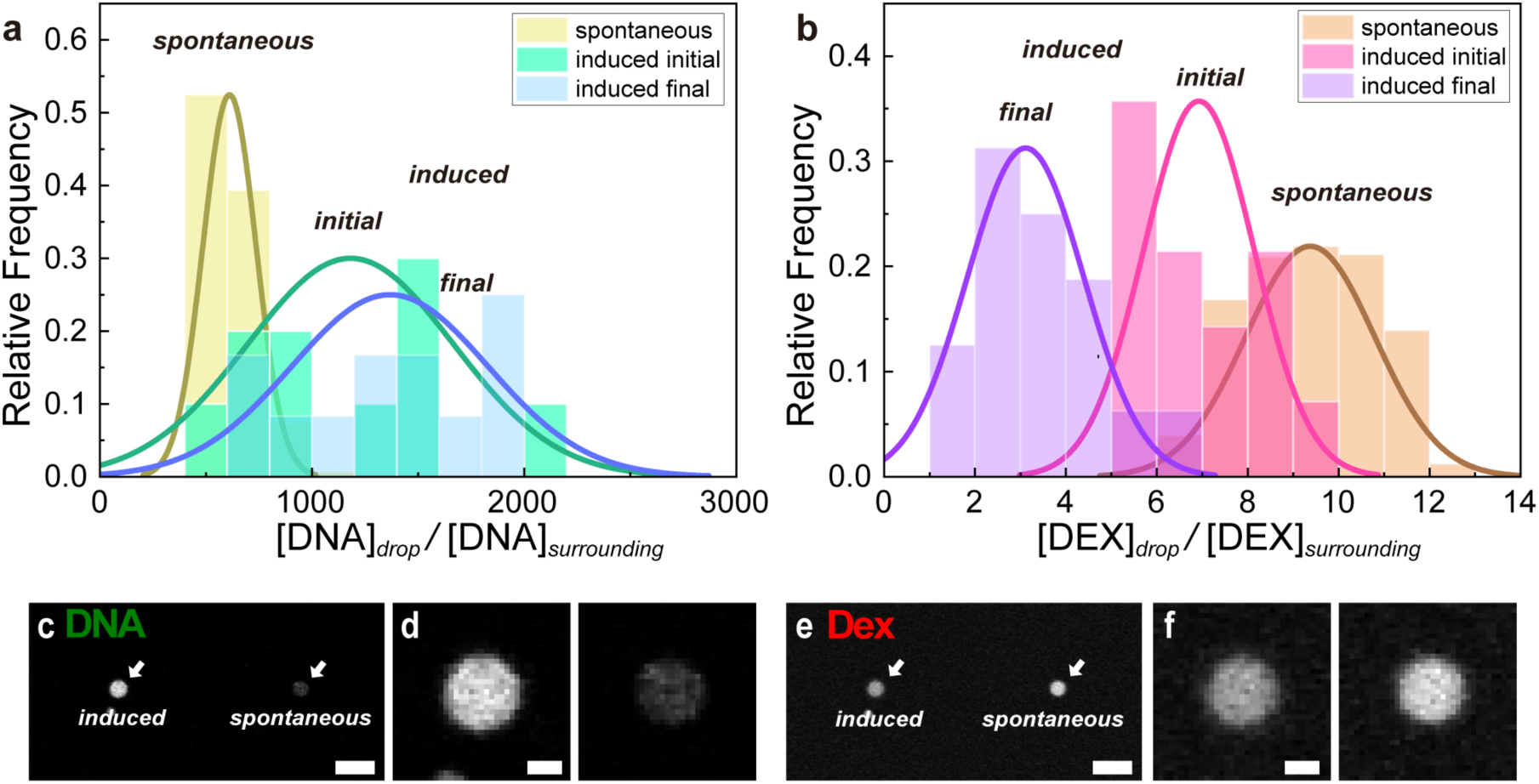
Comparison between LIPS droplets and the droplets by spontaneous phase separation. **a-b**, Histogram of the DNA enrichment factor and DEX concentration ratio of LIPS droplets and spontaneous droplets. The initial (*t* = 0 h) and the final (long-time) values for LIPS droplets are displayed. The LIPS droplets were generated from the upper PEG-rich phase of DEX 6 wt.% and PEG 2.6 wt.% with DNA (*λ*-DNA (48,502 bp) 29 ng/µL). Spontaneous droplets were produced from a two-phase mixture that has the same composition as the one used for the preparation of the upper PEG-rich phase for LIPS experiments. **(a)**; Histogram of the DNA enrichment factor. (**b)**; Histogram of concentration ratio of DEX between the droplet and the surrounding phase. **c-f**, Confocal images of a LIPS droplet generated by laser irradiation for 7 min and a droplet generated by spontaneous phase separation. The droplets were observed at *t* = 7 min after switching off the laser. **(c)**; Confocal images of DNA. (left) LIPS droplet. (right) Droplet obtained by spontaneous phase separation. Scale bar = 10 μm. **(d)**; Magnified image of droplets in Fig. 5c. Scale bar = 2 μm. **e**, Confocal image of DEX. Scale bar = 10 μm. **(f)**; Magnified image of droplets in Fig. 5e. Scale bar = 2 μm.

For the direct comparison of LIPS droplets and spontaneous droplets, we conducted LIPS with ATPS of DEX/PEG; the PEG-rich phase solution containing spontaneous DEX-rich droplets (Fig. 5c-f). LIPS droplets were generated as same as in the other experiments. The fluorescent intensity of DNA in the LIPS droplet at *t* = 7 min after generation was higher than the spontaneous one by a factor of 4.3. The intensity of DEX was lower than the spontaneous one by a factor of 0.7. Thus, the direct comparison confirmed that LIPS droplets have higher *EF*^DNA^ and lower *CR*^DEX^ than those of spontaneous droplets. These observations indicate that the property of LIPS droplets is not very far from that of spontaneous phase separation, but different from it. In the PEG concentration in this study, the Soret effect of DNA itself(33) could contribute to the mechanism for the higher *EF*^DNA^ for LIPS droplets than that of spontaneous one(34, 35). If so, *EF*^DNA^ could increase further by increasing *t*_IR_ up to the equilibrium of the Soret effect. Braun et al. demonstrated laser-induced DNA enrichment using the Soret effect and convection although it is not a phase separating system(36). The enriched DNA in their method diffused away after heating. In our case, convection is negligible since local heating is achieved by heating the ITO surface at the bottom of the sample cell. Note that we confirmed that the Soret effect of DNA itself is not the mechanism for the stable DNA enrichment in LIPS droplets (*SI Appendix*, section 7).

## Composition of LIPS droplet

The composition of LIPS droplets was estimated in confocal measurements by spiking fluorescently labelled DEX and PEG in DEX/PEG mix (See Methods and *SI Appendix*, Fig. S5). The initial and final DEX/PEG compositions of LIPS are plotted as the dotted and closed blue circles in *SI Appendix*, Fig. S5d, respectively. These compositions were estimated from the mean values of *CR*^DEX^ and *CR*^PEG^ at the initial state, i.e., just after LIPS: *t* ∼ 5 min and after long-time incubation: *t* ∼ 100 min (*SI Appendix*, Fig. S5a, b, c). The initial composition is close to the composition of the DEX-rich phase in the spontaneous ATPS of DEX/PEG (red triangle in *SI Appendix*, Fig. S5d), while the final composition is close to the PEG-rich phase (red circle in *SI Appendix*, Fig. S5d). This means that the DEX/PEG composition of LIPS droplet gradually changes with time, approaching to the composition of PEG-rich phase. However, as seen in the stable plateau of *CR*^DEX^ in Fig. 4c, the LIPS droplet almost pauses the composition change to stay at a particular composition in the metastable state.

## Patterning of LIPS droplets

Finally, we attempted to demonstrate the controllability, reproducibility, and longevity of LIPS droplets by patterning LIPS droplets in the presence of DNA along a designated structure. Multiple LIPS droplets were generated at the defined position to draw a heart mark by laser irradiation for 5 min (*SI Appendix*, Movie 2). The time interval between each LIPS was 1 min and the distance between droplets was approximately 50 μm. Fig. 6 shows the fluorescence image of DNA-containing LIPS droplets, stained with the intercalator dye. The patterned LIPS droplets demonstrate not only the controllability of LIPS but also that LIPS droplets do not interfere with the formation of neighbor droplets under the present condition. More complicated patterning should be possible if our method is combined with laser foci technique (37, 38). In preliminary experiments, we also found that it is possible to translocate LIPS droplets by moving the laser focal point. Exploiting these phenomena, the coalescence of two droplets was also demonstrated via manipulation (*SI Appendix*, Movie 3). This demonstration confirms the fluidity of the LIPS droplets, stating that the longevity is not caused by trivial processes such as gelation or other chemical reactions. We also observe the abrupt disappearance of LIPS droplets by short laser irradiation (*SI Appendix*, Movie 4). Thus, the current system allows for the microscopic handling of LIPS droplets: the patterning, translocation, fusion, and clearance of the DEX-rich droplets, although the precise understanding of the mechanism for these phenomena needs further studies.

**Fig. 6:**
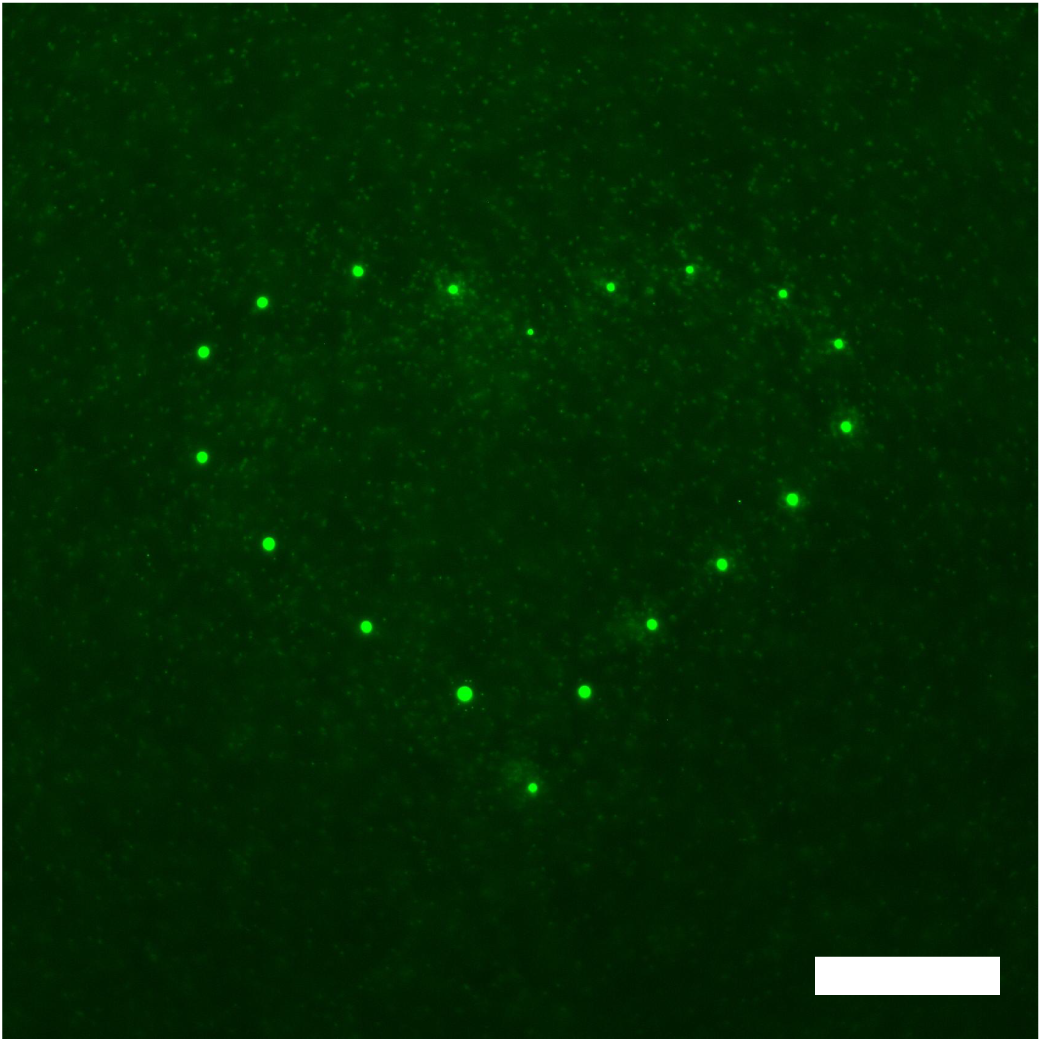
Heart shape patterned by DNA-enriched droplets. Fluorescent image of *λ*-DNA (48,502 bp) observed by epifluorescence microscopy. Each droplet was generated by the laser irradiation of the sample for 5 min to the upper PEG-rich phase of DEX 6 wt.% and PEG 2.6 wt.% + *λ*-DNA (48,502 bp) 29 ng/µL. Scale bar = 100 μm.

## Discussion

The most prominent finding of this study is the extraordinarily long-time stability of the LIPS droplets which allows stable DNA enrichment. The generated droplets in our method are principally non-equilibrium droplets because the initial state of the sample is a homogeneous single PEG-rich phase. Intuitively, the droplets were expected to disappear, as observed in the LIPS experiments conducted thus far(21, 25). However, the generated droplets did not vanish for a significantly long time after the laser was disabled. We observed a distinct decrease in the diameter, indicating that the system moved towards the original point on the coexisting curve. However, the shrinkage stopped at a point before returning to the original homogenous PEG-rich phase. Resultantly, the DEX/PEG composition of the droplets was close to but not exactly on the phase boundary (*SI Appendix*, Fig. S5). These observations mean that LIPS droplets were stacked in a metastable state. The stable LIPS droplets were also generated by using ATPS of DEX/PEG with different compositions, suggesting the generality of the stable LIPS droplets. Regarding the metastability, it is also worth noting the laser-induced disappearance of LIPS droplets (*SI Appendix*, Movie 4). The laser irradiation seems to work as a perturbation to agitate the metastable droplets and the surrounding phase, and as a result, the droplets returned into the most stable state of the homogeneous phase. Consistent with this interpretation, spontaneously formed DEX-rich droplets in DEX/PEG ATPS which are stable in nature were retained even after the laser irradiation (*SI Appendix*, Movie 5).

The initial condition of the sample on the phase boundary would be a key to the mechanism for the longevity of LIPS droplets. For the system exactly on the phase boundary, the other coexisting phase is energetically the same. This implies that, in principle, the system can shift towards both directions. This can be a similar situation to the coexistence of the ice at 0 ℃ in water at 0 ℃. There should be competing effects between two equilibrium phases to which the system should go. The initial speed of droplet shrinkage decreased when the system was approaching the coexisting curve (Fig. 2f), suggesting that the driving force to cross the phase boundary to return toward the original equilibrium in the one-phase region dramatically decreases and this causes the longevity as schematically illustrated in Fig. 2g and h. On the other hand, this contention does not give full explanation for the reason why the system has metastable states.

The present study indicates that there is an unknown mechanism which prevents the system from crossing the phase boundary to return to the one-phase region. According to the classical nucleation theory for phase separation, there is a critical radius *R*_c_ for droplet growth, for which the energy barrier is determined by the interfacial energy and the bulk free energy of a droplet. While droplets of radius smaller than *R*_c_ preferentially shrink, droplets show a stable growth when they occasionally reached *R*_c_(39). In the reverse process, it should be different from the same *R*_c_, but there would be a similar energy barrier for droplet shrinkage when the system crosses the phase boundary from the two-phase region. That might be a mechanism for the remarkable longevity of the LIPS droplets. Of course, this model would be too simplified. The exact understanding of the molecular mechanism for the metastability of LIPS droplets needs future study.

In recent years, many studies reported that intracellular droplets of biomolecules, termed ‘cell biological droplets’ play pivotal roles in biology(1–3, 40–42). Thereby, the physicochemistry of phase separation systems attracts large attention not only in soft matter physics but also in chemical biology, biophysics and molecular biology(43–45). The metastable droplet generation mechanism in this study might be related to the formation of stable cellular structures in protocells. It would be interesting to apply our method to systems consisting of cell extract or living cell systems. The DEX/PEG system investigated herein also attracts attention because the DEX-rich droplet is a model system to build protocell or artificial cell systems(6). Our method can be utilized to produce cellular structures in artificial cells or to the local formation of DNA origami structures. So far, we could not keep both the LIPS droplet and the surrounding phase at the same temperature in principle, but the present method enables us to do it. Our method has no limitation in generation temperature and the temperature can be changed even after generation (*SI Appendix*, section 3), which provides a way to develop new tools for biochemical reactions and digital bioassays such as digital PCR. Further, the PEG-rich phase enables the enrichment of rare substances and their manipulation from its dilute solution, which could be applied to drug delivery. We preliminarily confirmed that RNA could also be enriched in the LIPS droplets. Proteins such as antibodies can be enriched in DEX-rich droplets, in particular when they are tagged with dextran-binding domain (DBD)(8). Considering these, the present LIPS method would be a technological basis for the microscopic manipulation of biological molecules/systems by use of LIPS droplets as a carrier.

In summary, we succeeded in metastable LIPS droplet generation with unexpected longevity which has never been observed before. The counterintuitive longevity is putting a fundamental question to physics. It is also the first report of success in LIPS DEX droplet generation using the DEX/PEG system and in LIPS DNA droplet generation which keeps fluidity. We achieved highly stable DNA enrichment by a factor of thousands. The advantage of our system is its applicability to a variety of biomolecules not only DNA and RNA but also proteins such as enzymes and antibodies tagged with dextran-binding domain (DBD). We conducted the concentration-driven LIPS using the Soret effect, which gave controllability of generation temperature. Simple setup by conventional equipment gave accessibility for laboratory use. Those features widen the possibility of applications of LIPS droplets in biological research. The properties of encapsulating substances accompanied by phase separation, are important also in pharmaceuticals(46), medical science, and agricultural science.

## Materials and Methods

### Materials

We prepared a sample by mixing Dextran (DEX, Mw = 4.5 – 6.5 × 10^5^, Sigma), Polyethylene glycol (PEG, Mn = 3.5 × 10^4^, Aldrich), λ-DNA (48,502 bp, Nippon Gene), and pure water. To visualize and evaluate the concentration of polymers, we used TRITC-Dextran (Mw = 5.0 × 10^5^, Sigma) and FITC-PEG (Mw = 3.0 × 10^4^, Creative PEG Works). DNA was visualized using an intercalator (SYBR Gold; Invitrogen). DEX and PEG were dissolved in water at 20 wt.% as stock solutions before mixing.

The phase diagram of the system is shown in Fig. 1a. We chose the system of the above molecular weights of DEX and PEG because the temperature dependence of the phase diagram is negligible. This means that our droplet generation is not induced by temperature-driven phase separation but by the Soret effect. We confirmed this property experimentally by the fact that phase separation cannot be induced by heating the bulk sample (see also *SI Appendix*, section 2). Note that the temperature effect to phase separation in DEX/PEG system depends on the molecular weight of polymers and there are other DEX/PEG systems which show clear temperature dependence for the lower molecular weight of DEX and PEG(28, 47).

For the droplet-generation experiments, we first prepared a sample with an initial composition in the two-phase region to experimentally obtain a sample on the coexisting curve. All components were mechanically mixed using a conventional vortex mixer and centrifuged at 15,000 rpm (Eppendorf Himac Technologies, CF18RS relative centrifugal force of 16,451.37*g*) for 20 min under temperature control at 298 K. The upper PEG-rich phase (indicated by an arrow in Fig. 1a) produced from the mixture was extracted and used as a sample. For the mixture of DEX 2.6 wt.% PEG 6 wt.% (open circle in Fig. 1a), the composition of two separated phases after centrifugation estimated from the fluorescent intensity of a dye (TRITC-DEX, FITC-PEG) using a microplate reader (Molecular Devices, SpectraMax iD3) was DEX 0.7 wt.% and PEG 3.1wt.% for the upper PEG-rich phase and DEX 8.4 wt.% and PEG 0.9 wt.% for the lower DEX-rich phase, where we assumed that the concentration of the polymer with the dye was directly proportional to that of the dye-free polymer of the same type. The quantity of fluorescently labelled DEX varied from 0.02%–0.2% depending on the experiment. The concentrations in the text denote the total DEX concentration. The DNA concentration of each equilibrated phase was 70 pg/μL for the upper PEG-rich phase estimated by the confocal image of DNA, and 60 ng/μL for the lower DEX-rich phase estimated by spectroscopy (Thermo Scientific, NanoDrop 2000).

To observe the spontaneously forming droplets, we used a vortex mixer to agitate the solution for a few minutes. Subsequently, we used the upper PEG-rich phase containing DEX-rich droplets after waiting for a sufficient amount of time for sedimentation to reduce the number of droplets for 30 min.

### Local heating experiments and microscopy

Local heating was performed under a microscope (Nikon Ti2E). An infrared (IR) laser with a wavelength of 1,064 nm was connected to the microscope and focused on the sample (Thorlabs; Diode laser 1064 nm, a laser power of 30 mW at the sample position). The sample was sandwiched between a cover glass and a glass plate with ITO (Indium Tin Oxide) coating (EHC. Co. Ltd.; ITO thickness = 20*–*40 nm, and resistance = 100 ohms). ITO absorbs light of wavelength 1,064 nm, which causes a temperature increase at the beam position. The sample thickness was controlled by a polyimide film with a thickness of 85 *µ*m. The temperature of the sample cell was controlled at 298 ± 0.1 K by a temperature stage (Linkam PE120) with a Peltier module. For multiple-droplet generation, temperature control is necessary to prevent the temperature gradient caused by laser irradiation from influencing the droplets. The maximum temperature increase by local heating was approximately 10 K, which was estimated from the temperature dependence of fluorescence intensity of Rhodamine B solution in water (*SI Appendix*, section 1). The microscope images were captured using a CMOS camera (Andor Zyla).

For confocal microscopy, the droplets were generated using the local heating setup described above and they were observed under a confocal microscope (Leica Microsystems, SP-8). The glass plate (thickness of 0.7 mm) with an ITO coating is greater than the working distance of the objective lens (100X, N.A. 1.47) used in confocal microscopy. The sample cell was flipped, and the induced droplets were observed from the bottom. For the simultaneous observation of induced and spontaneous droplets, a glass plate (thickness 0.2 mm) with an ITO coating was used without flipping over the plate. A white laser was used for excitation (488 nm for SYBR gold or FITC-PEG and 561 nm for TRITC-DEX), and the experiment was performed with the minimum number of acquisitions to reduce the contribution of photobleaching. We did not perform 3D acquisition every time in a time-course measurement and analyzed kinetics using the droplet diameter instead of droplet volume for the same reason. Under this condition, the decrease in intensity caused by photobleaching can be considered negligible (*SI Appendix*, section 5). The background light caused by the reflection of the laser from the substrate was removed by notch filters and time-gated acquisition from 0.2 ns. The time evolution was observed at room temperature (297 ± 0.5 K). The droplet diameter was estimated by contour detection using commercial software (Image Pro Plus, Media Cybernetics). As the obtained values could be underestimated, we set the error bar to ± 0.5 μm. For the non-linear fit analysis of the droplet diameter, we added the initial droplet diameter after LIPS (*t* = 0) determined from the phase contrast microscopy image to the time evolution of the droplet diameter observed by confocal microscopy as the value *d*_0_ at *t* = 0.

For the estimation of the composition of the LIPS droplets, we performed the LIPS droplet generation experiments on the sample including fluorescently labelled DEX and PEG (TRITC-Dextran (Mw = 5.0 × 10^5^, Sigma) and FITC-PEG (Mw = 3.0 × 10^4^, Creative PEG Works)). The concentration ratios of DEX (*CR*^DEX^) and PEG (*CR*^PEG^) between the droplet and the surrounding phase were determined as a fluorescent intensity ratio. They were converted to the concentration by multiplying the concentration of the surrounding phase (0.7 wt.% for DEX, 3.1 wt.% for PEG).

### Estimation of absolute DNA concentration

We determined the DNA concentration by comparison with the confocal data of DNA suspension in water with SYBR gold (See also *SI Appendix*, section 4). At a low concentration range below 100 pg/μL, DNA molecules are isolated and visible in confocal images because the full length of λ-DNA (48,502 bp) is approximately 16 μm. We counted the number of DNA molecules in several images and obtained the concentration dependence of the number per unit area (*SI Appendix*, Fig. S10). At the concentration range above 100 pg/μL, it becomes difficult to distinguish the isolated DNA molecules. We observed the confocal image of DNA suspensions in the same sample cell and other experimental conditions with that of the LIPS experiments. The concentration dependence of the average fluorescent intensity was measured and used to determine the absolute concentration of the LIPS droplets.

## Acknowledgements

The authors thank Masao Doi for stimulating discussions on the mechanism of metastability. They thank Hajime Tanaka for helpful discussions on phase separation kinetics and his constant encouragement. They are grateful to Takahiro Muraoka for valuable discussions and his encouragement. They also thank Yutaro Ii for providing RNA samples, Shinichiro Kanatsuki for technical support of temperature control, and Kazutaka Irisawa for the optimization of ITO coating. This work was supported in part by Grant-in-Aid for Scientific Research (S; JP19H05624 to H.N.) and (C; 19K03763 to M.K.) from the Japan Society for the Promotion of Science (JSPS), and JST CREST, Japan (JPMJCR19S4 to H.N.), and the Kao Foundation for Arts and Sciences (Kao Crescent award to M.K.).

## Author contributions

H.N. designed the project and supervised the study. M.K. performed the experiments and discovered the long-time presence of LIPS droplets. M.K. interpreted the physical mechanism underlying the phenomenon and analyzed the data. All authors discussed the results. M. K. and H. N. prepared the manuscript.

