## Supplementary Information for "Metastable phase-separated droplet generation and long-time DNA enrichment by laser-induced Soret effect"

*Tokyo 113-8656, Japan*

**Contents**

**Supplementary Movie 1: Droplet generation process by LIPS**

**Supplementary Movie 2: Heart shape patterning process of LIPS droplets**

**Supplementary Movie 3: Coalescence of two LIPS droplets**

**Supplementary Movie 4: Laser-induced disappearance of LIPS droplet**

**Supplementary Movie 5: Laser irradiation to a spontaneous droplet**

**Supplementary Figures S1 to S20**

**Supplementary Text 1: Temperature increase by the laser irradiation**

**Supplementary Text 2: Case without ITO coating**

**Supplementary Text 3: Heating of a LIPS droplet**

**Supplementary Text 4: Estimation of DNA enrichment factor and Dex concentration ratio**

**Supplementary Text 5: Validity of the temporal change of the intensity ratio**

**Supplementary Text 6: DNA enrichment factor and DEX concentration ratio of droplets by. spontaneous phase separation**

**Supplementary Text 7: Localization of DNA in a PEG solution**

**Movie 1: Droplet generation process by LIPS**

The laser irradiation process for 10 min and the scenario after disabling the laser. This process was observed using phase contrast microscopy. Sample: upper phase of DEX (Mw550,000) 6 wt.% + PEG (Mw35,000) 2.6 wt.% + λ-DNA 29 ng/μL. A square with a side of 169 μm. Playback speed 10X.

**Supplementary Movie 2: Heart shape patterning process of LIPS droplets**

Patterning process of LIPS droplets in a heart shape. Sample: upper phase of DEX (Mw550,000) 6 wt.% + PEG (Mw35,000) 2.6 wt.% + λ-DNA 29 ng/μL. A square with a side of 676 μm. Playback speed 100X.

**Supplementary Movie 3: Coalescence of two LIPS droplets**

Two DEX droplets generated by LIPS were coalesced by manipulation. The process was observed by phase contrast microscopy. Sample: upper phase of DEX (Mw550,000) 6 wt.% + PEG (Mw35,000) 2.6 wt.%. A square with a side of 84.5 μm. Playback speed 1X.

**Supplementary Movie 4: Laser-induced disappearance of LIPS droplet**

The laser irradiation process to the LIPS droplet observed by phase contrast microscopy. The droplet disappeared after successive laser irradiation. The final disappearance depends on the balance between the mixing effect and the induction of new droplets by laser irradiation. Sample: upper phase of DEX (Mw550,000) 6 wt.% + PEG (Mw35,000) 2.6 wt.%. A square with a side of 33 μm. Playback speed 5X.

**Supplementary Movie 5: Laser irradiation to a spontaneous droplet**

The laser irradiation process to a spontaneous droplet observed by phase contrast microscopy. The droplet did not disappear after successive laser irradiation. The image contrast becomes low during the laser irradiation, but the droplet still appears after switching off the laser. Sample: upper phase of DEX (Mw550,000) 6 wt.% + PEG (Mw35,000) 2.6 wt.%. A square with a side of 33 μm. Playback speed 5X.

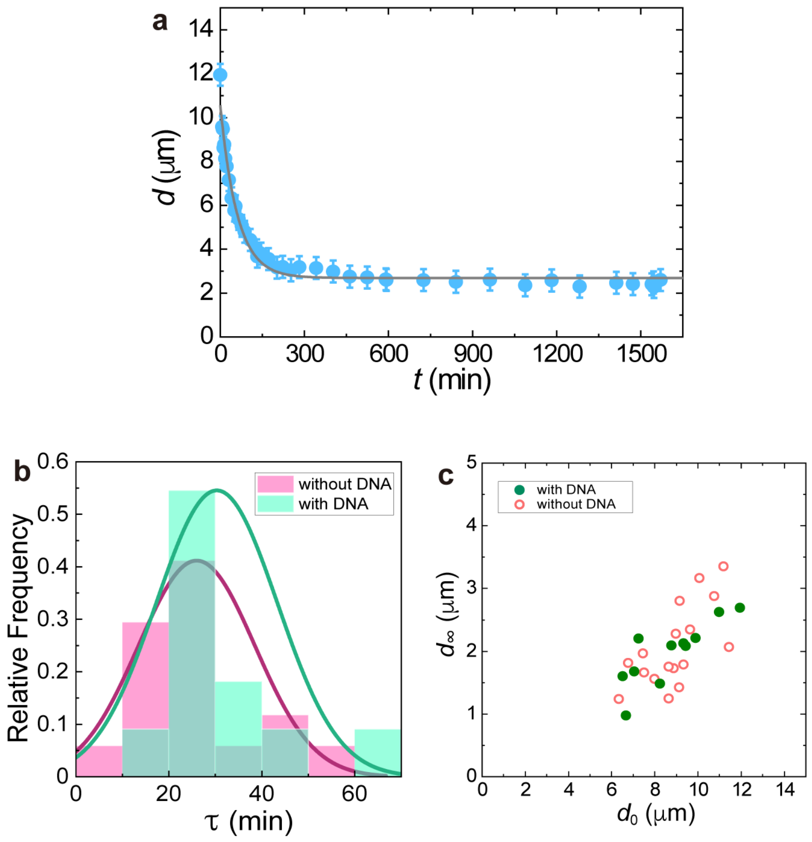


**Fig. S1: Time evolution of droplet diameter. a**, Time evolution of the droplet diameter for the same experiment shown in Fig. 4a-c in the main text. The droplet was generated by laser irradiation for 12 min to the upper PEG-rich phase of DEX 6 wt.% and PEG 2.6 wt.% + $\lambda$-DNA (48,502 bp) 29 ng/μL. The value at $t = 0$ was obtained from the phase-contrast image. The rest were obtained from the confocal images of DNA. The diameter approaches a finite value. The solid curve was obtained by fitting the data to a function, $d(t) = (d_0 - d_\infty)\exp(-t/\tau) + d_\infty$, where $d_0$ and $d_\infty$ represent the diameters at $t = 0$ and $t = \infty$, respectively, and $\tau$ represents the characteristic decay time. The fitting parameters from the fit were $\tau = 60.7$ min and $d_\infty = 2.7$ μm. **b**, Histogram of the decay time of droplet diameter in the case with and without DNA. **c**, Final droplet diameter $d_\infty$ at $t = \infty$ estimated by the fit plotted against the initial droplet diameter $d_0$ at $t = 0$ (measured from phase contrast image). Green circles: cases with DNA. Red open circles: without DNA. The droplets were generated from the upper PEG-rich phase of DEX 6 wt.% and PEG 2.6 wt.% with/without DNA ($\lambda$-DNA (48,502 bp) 29 ng/μL)).

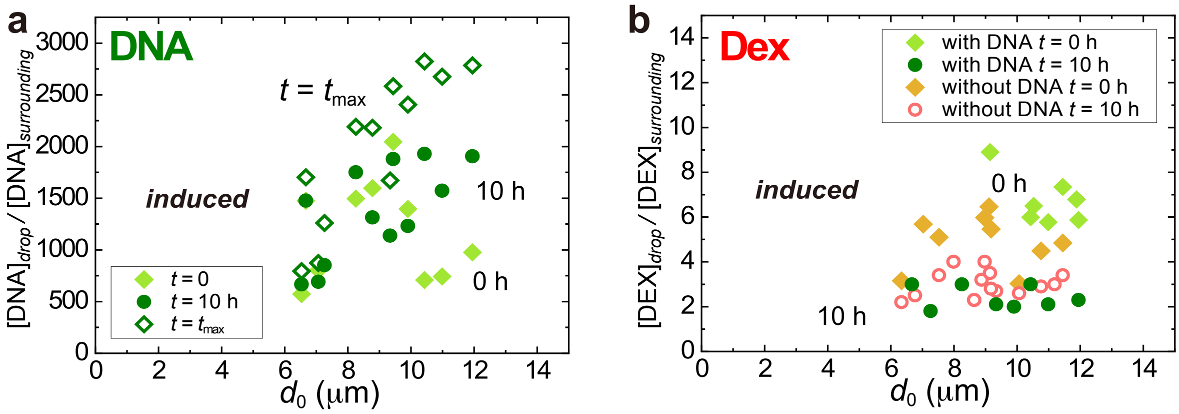


**Fig. S2: The DNA enrichment factor and DEX concentration ratio. a**, The DNA enrichment factor of LIPS droplets. The initial value ($t$ = 0 h), maximum value ($t = t_{max}$), and long-time value ($t$ = 10 h). The droplets were generated from the upper PEG-rich phase of DEX 6 wt.% and PEG 2.6 wt.% with DNA ($\lambda$-DNA (48,502 bp) 29 ng/μL). **b**, Concentration ratio of DEX between the droplet and surrounding phase of the induced droplets at $t$ = 0, 10 h estimated from the fluorescence intensity ratio of dextran. The droplets were generated from the upper PEG-rich phase with/without DNA (DEX 6 wt.% and PEG 2.6 wt.% with/without DNA ($\lambda$-DNA (48,502 bp) 29 ng/μL).

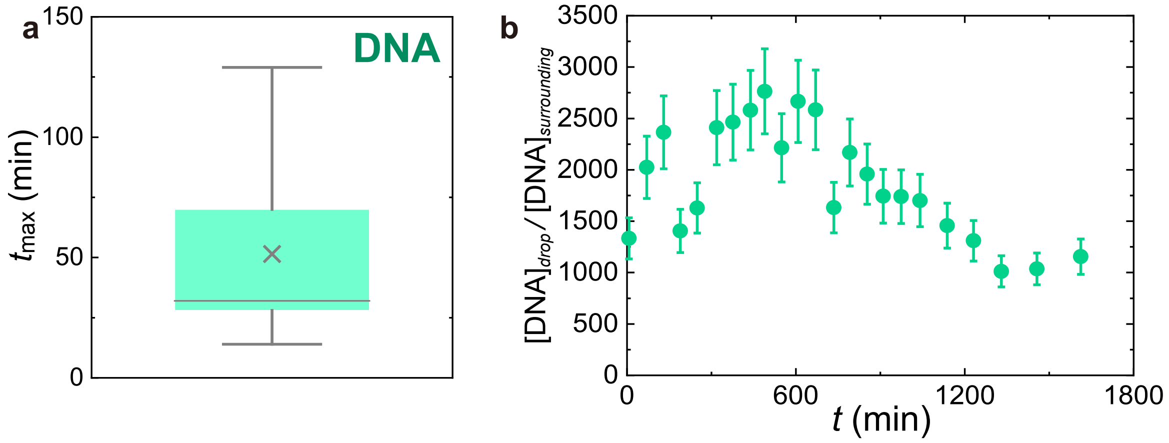


**Fig. S3: Behavior of DNA enrichment factor to reach its maximum value. a**, Time scale when the enrichment factor of DNA ($EF^{DNA}$) reached its maximum value. 25–75 percentile (box), median (line), mean (cross), highest/lowest observations without outliers (whiskers). **b,** A case where $EF^{DNA}$ showed a broad time change and showed its maximum value at a very long time around 500 min. The factor $EF^{DNA}$ was kept at a high value of more than 2500 for an exceedingly long time.

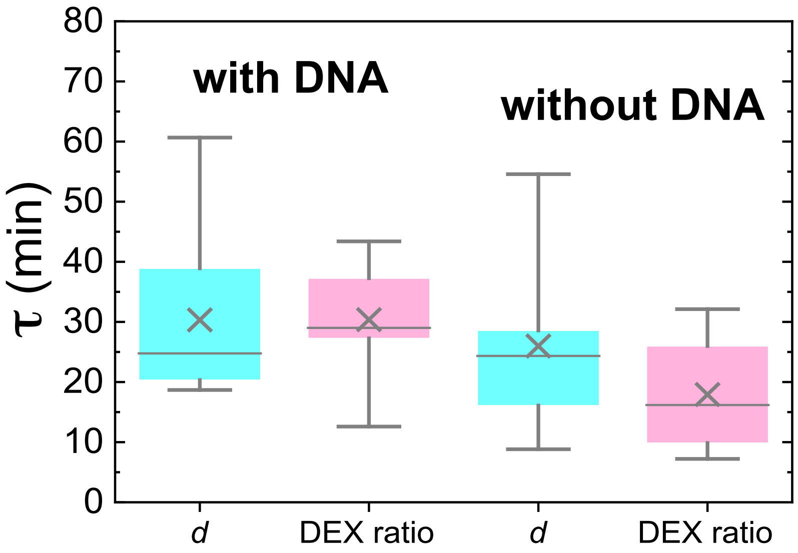


**Fig. S4: Comparison of timescale between the droplet shrinkage and the DEX concentration change.** Decay time was estimated by exponential fit. The function used for the fit was the same expression as that given in Fig. S1. Decay time of droplet diameter (light blue) and DEX concentration ratio (pink) in the case of w/ and w/o DNA. 25–75 percentile (box), median (line), mean (cross), highest/lowest observations (whiskers).

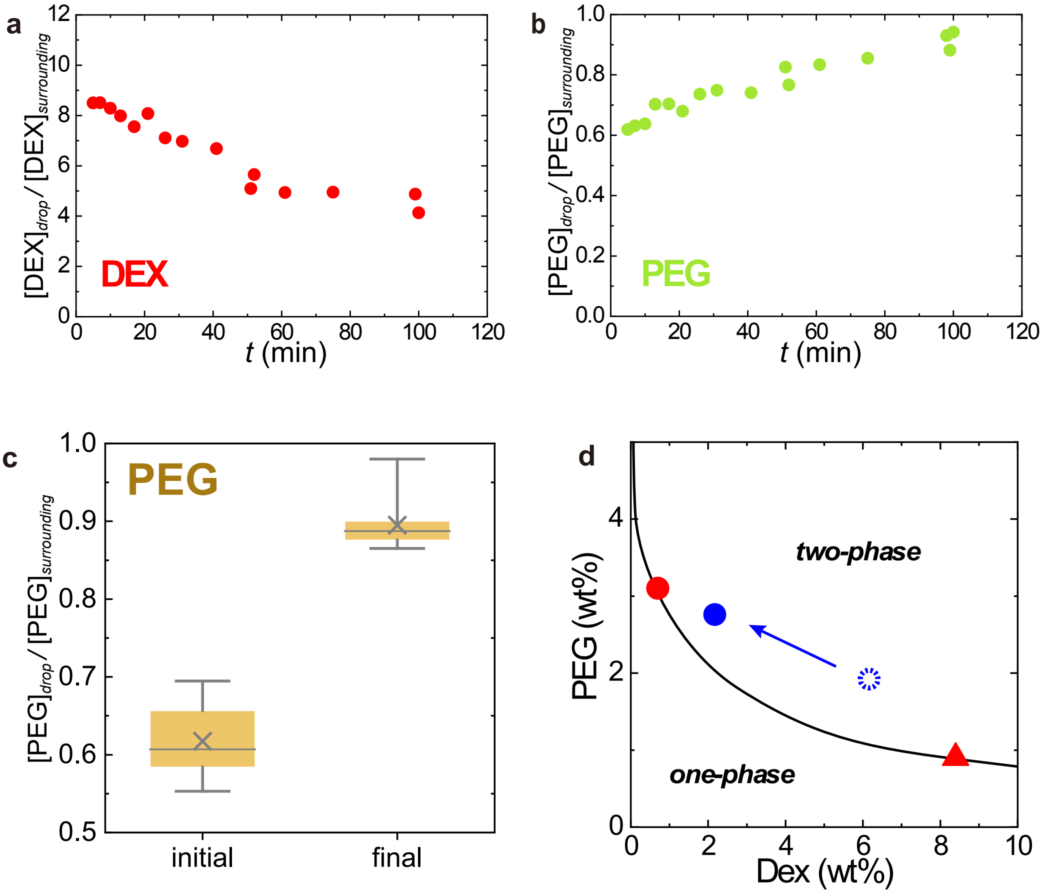


**Fig. S5: Observation of the composition of LIPS droplets. a-b**, Typical time evolution of (**a**) DEX and (**b**) PEG concentration ratios of a LIPS droplet. The droplet was generated by laser irradiation for 11 min to the upper PEG-rich phase of DEX 6 wt.% and PEG 2.6 wt.% + $\lambda$-DNA (48,502 bp) 29 ng/µL (including 0.2 wt% of TRITC-Dextran and FITC-PEG). The concentration ratios of DEX and PEG were estimated from the fluorescent intensity of dye (TRITC-DEX and FITC-PEG) in confocal measurements. **c**, PEG concentration ratio. 25–75 percentile (box), median (line), mean (cross), highest/lowest observations (whiskers). Data w/ and w/o DNA are both included because there was no clear difference beyond experimental error. **d**, The final composition of the LIPS droplets in a metastable state which is shown in the phase diagram. The final composition (closed blue circle) was estimated from the long-time value of DEX and PEG concentration ratio of the LIPS droplet by confocal measurements. The composition just after generation (at around $t$ = 5 min) is indicated by a dotted circle. Red circle: upper PEG-rich phase used for LIPS experiment, red triangle: lower DEX-rich phase. The solid curve is the coexisting curve. The composition changed with time approaching the original upper phase and stopped at the blue circle.

**Supplementary Text 1: Temperature increase by the laser irradiation**

We estimated the temperature increase attributed to laser irradiation from the temperature dependence of fluorescence intensity. The temperature dependence of fluorescence intensity of 0.01 wt.% aqueous solution of Rhodamine B is illustrated in Fig. S6a. The intensity was normalized by that at 25 °C, which was the temperature in the droplet experiment. The intensity monotonically decreased with an increase in the temperature. Figure S6b shows the intensity profile of a fluorescent image of the 0.01 wt.% Rhodamine B aqueous solution in a sample cell with ITO coating, which is captured after laser irradiation for 1 min. The intensity is normalized by the image captured before irradiation. The intensity at the laser-irradiation spot decreased from 1 to 0.85. The increase in temperature caused by laser irradiation was estimated at approximately 10 K. Changes in concentration attributed to the Soret effect of Rhodamine B are negligible (in the order of $10^{-5}$ wt.%/K).

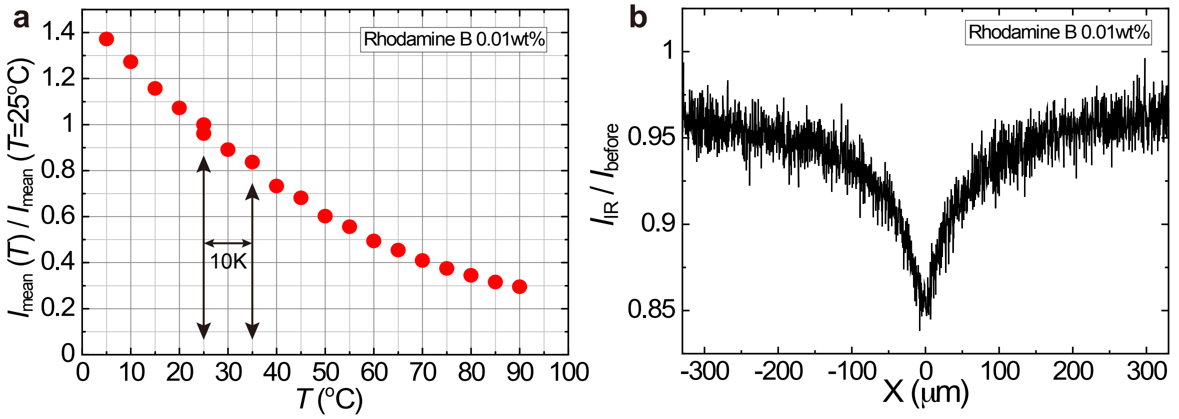


**Fig. S6: Estimation of temperature increase. a,** Temperature dependence of 0.01wt.% aqueous solution of Rhodamine B. **b,** Intensity profile of the fluorescent image of the sample after laser irradiation.

**Supplementary Text 2: Case without ITO coating**

We experimentally verified that our droplets were generated by the Soret effect by comparing two cases, i.e. with and without ITO coating (Fig. S7). When laser light is irradiated on the region without an ITO coating, no temperature gradient is produced. In this case, we did not identify any sign of a concentration change, which means the interaction between the sample and laser light does not induce phase separation in the sample and that local heating is essential for generating droplets.

Further, heating or cooling the bulk sample by a temperature stage in the range of 283–368 K did not cause anything to indicate a phase separation, as shown in Fig. S8. This confirms that the phase diagram of our system is inert to temperature. In other words, droplets cannot form without 'local' heating. Hence, the temperature gradient caused by local heating changes the local concentration due to the Soret effect, which results in phase separation.

The generation process (Fig. 1d in the main text) appears similar to that observed by Walton and Wynne(1, 2); however, the mechanisms are different. They estimated that the effect of thermophoresis is negligible in their system and concluded that their LIPS was caused by the electromagnetic energy stored in the sample by laser irradiation. In our case, the Soret effect causes droplet generation.

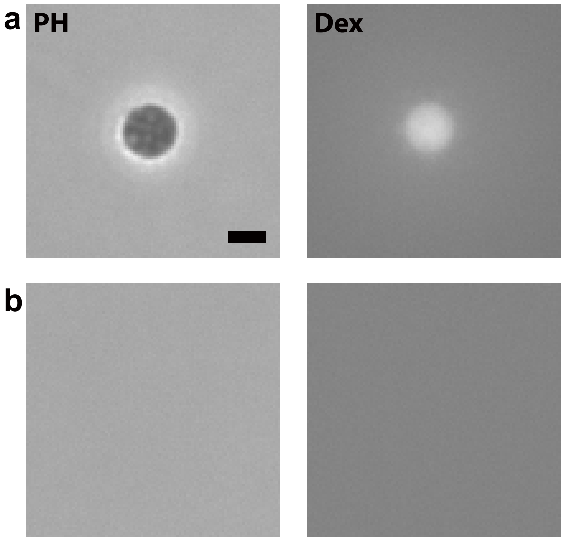


**Fig. S7: Comparison of two cases with/without ITO coating.** The droplet was induced by laser irradiation for 10 min to the upper PEG-rich phase of DEX 6 wt.% PEG 2.6wt.%. The same sample cell was irradiated with laser light in different regions (a) with and (b) without ITO coating. Left: phase-contrast images, right: fluorescent images of dextran. Scale bar = 5 μm.

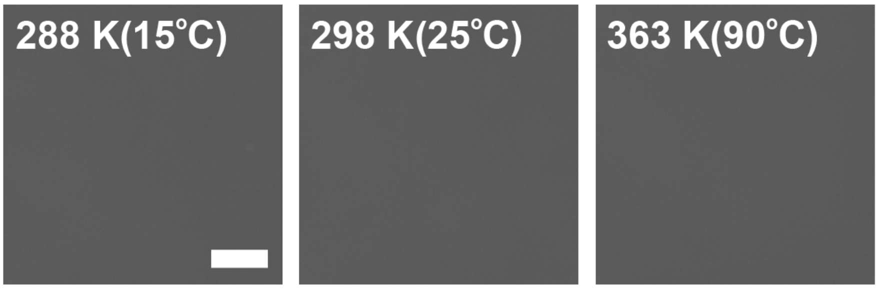


**Fig. S8: Temperature dependence of a bulk sample.** Phase contrast images of the upper PEG-rich phase of DEX 6 wt.% PEG 2.6wt.%. Scale bar = 20 μm.

**Supplementary Text 3: Heating of a LIPS droplet**

We examined if we can change the temperature of our LIPS droplet and it can be kept. We heated a LIPS droplet after switching off the laser, and the phase contrast images are shown in Fig. S9. The droplet is kept on heating or cooling. A decrease in the image contrast at high temperatures does not mean that the droplet is disappearing. The low contrast could be due to decreasing the difference in the refractive index between the droplet and the surrounding phase. The contrast returned to the original one when it was cooled again. This means that the droplet was stable on temperature change.

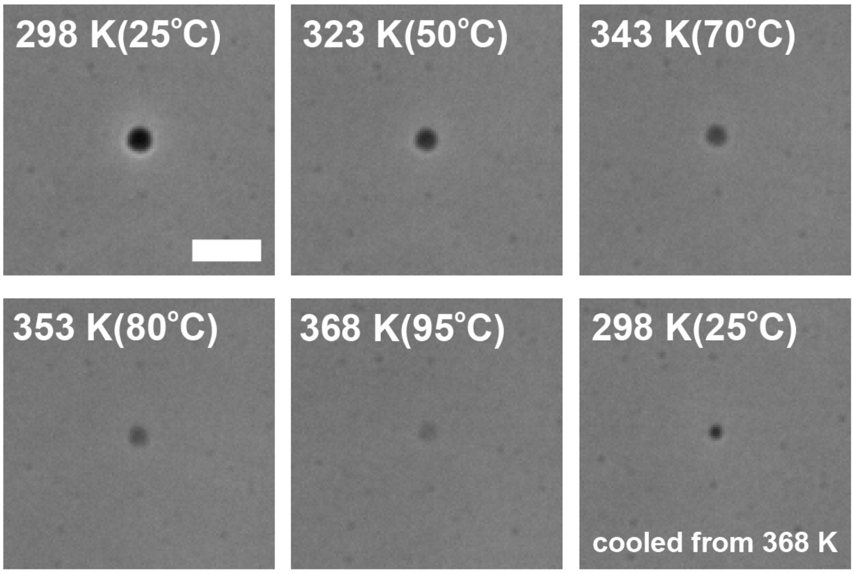


**Fig. S9: Droplet image on heating.** The droplet was first induced by laser irradiation for 5 min to the upper PEG-rich phase of DEX 6 wt.% PEG 2.6wt.% at 298 K. Phase-contrast images were taken when the whole sample cell was heated after generation. The image contrast of the droplet decreased on heating but returned on cooling to 298 K. Scale bar = 10 μm.

**Supplementary Text 4: Estimation of DNA enrichment factor and Dex concentration ratio**

**1. DNA enrichment factor**

We evaluated the DNA concentration of droplet $[DNA]_{droplet}$ and that of the surrounding phase (the upper PEG-rich phase) $[DNA]_{surrounding}$ separately, and we estimated the DNA enrichment factor as the concentration ratio $[DNA]_{droplet}$ / $[DNA]_{surrounding}$.

In the initial preparation of the DEX/PEG mix with DNA, we used 29 ng/μL of DNA as the concentration for the total mixture. However, most of it goes to the lower DEX-rich phase because DNA is enriched to the DEX-rich phase. Thus, the DNA concentration of the upper PEG-rich phase is very low (< 1 ng/μL). The concentration of nucleic acid below a few ng/μL is difficult to estimate by conventional methods such as UV spectroscopy. In our confocal experiments, the average intensity of fluorescence from SYBR gold is almost the same signal level as the one from pure water, implying that it is difficult to estimate accurate DNA concentration from the average intensity of fluorescent images. However, DNA molecules are visible because the full length of λ-DNA (48,502 bp) is about 16 μm. Thus, we estimated the low DNA concentration of the surrounding phase (upper PEG-rich phase) from the number of DNA molecules in a unit area of confocal images for calculating the enrichment factor.

For a low concentration range below 1 ng/μL, we observed DNA suspension in water by confocal microscopy at several DNA concentrations. The typical images are shown in Fig. S10a. We analyzed

10 images (image size: 155 μm × 155 μm) at each concentration and counted the number of DNA molecules in a unit area using commercial software (Image Pro Plus, Media Cybernetics). The spatial resolution of confocal measurements in the vertical (Z) direction was 193 nm (objective lens 100 X oil, N.A. 1.47), and the overlap of molecules in the Z-direction can be neglected. The DNA molecules were well isolated, and the analysis was successful below 600 pg/μL. The results are illustrated in Fig. S10b. The concentration dependence of the number of DNA molecules showed a good linearity in the concentration range of 3–600 pg/μL. We performed a linear fit to the concentration dependence and the result was used to estimate the DNA concentration of the upper PEG-rich phase used for LIPS experiments. The confocal images of the upper PEG-rich phase were analyzed and estimated as $[DNA]_{surrounding} = 70 \pm 10$ pg/μL. A confocal image of the upper phase is shown in Fig. S10c.

For the high concentration range above 1 ng/μL, we evaluated the DNA concentration from the absolute fluorescent intensity of confocal images. We observed DNA suspension in water in the same cell used in the LIPS experiments and measured the average intensity of the image at the same Z-position of droplet generation close to the ITO surface. The result is shown in Fig. S11. The linearity is maintained in the concentration range of 1–1000 ng/μL. We increased the amount of SYBR gold concentrate at high concentrations above 100 ng/μL to maintain a sufficient amount of SYBR gold for DNA molecules. For the suspension of 100 ng/μL, the intensity was measured in both conditions at 1/1000 and 1/100 of SYBR gold concentrate and the intensities were consistent with each other. In the LIPS experiment, we prepared all samples with 1/1000 of SYBR gold concentrate. The amount of SYBR gold is sufficient because the total DNA concentration for the initial mixture is 29 ng/μL, which is in the linear regime in the concentration dependence shown in Fig. S11. In addition, the initial DNA concentration of the PEG-rich phase was approximately 70 pg/μL, as shown above. The volume of one droplet was estimated to be in the order of 10 $\mu m^3$. This value is significantly smaller than the total sample volume by a factor of $10^{-8}$. We concluded that the sample contains an adequate amount of DNA molecules and SYBR gold to achieve high concentration in the droplet, and that linearity is maintained under the experimental conditions of this study.

Figure S12a shows an example of the time evolution of the absolute intensity of the LIPS droplet obtained as raw data. We converted the absolute intensity to DNA concentration (Fig. S12b). The enrichment factor was obtained by dividing the DNA concentration of the droplet by that of the surrounding phase (upper PEG-rich phase), i.e. $[DNA]_{droplet}$ /$[DNA]_{surrounding}$ (Fig. S12c). The experimental error of the enrichment factor was attributed to the estimation of the concentration value of the surrounding phase (upper PEG-rich phase) and the focal position in the confocal measurements. We considered these factors and determined the error bars in Fig. 12b in the main text as 15%.

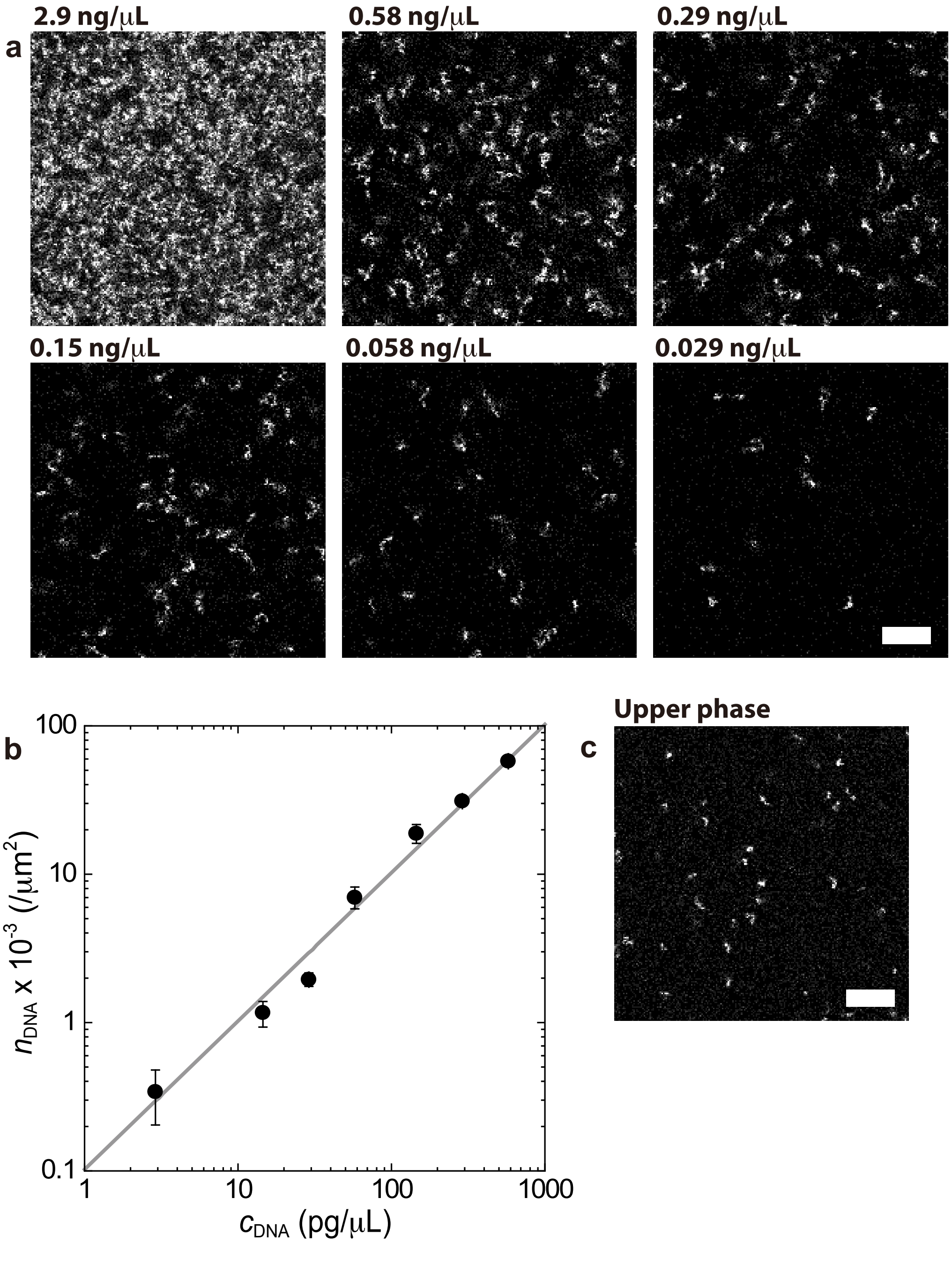


**Extended Data Fig. S10: Confocal images of DNA suspensions. a**, DNA suspension in water at several concentrations, $\lambda$-DNA (48,502 bp) visualized using DNA intercalator (SYBR gold). These samples were observed in a sample cell made of two coverslips with an 85-μm-thick spacer. The experiments were performed at room temperature (297 ± 0.5 K). Scale bar = 10 μm. **b**, Number of DNA molecules per unit area as a function of DNA concentration observed in DNA suspensions in water. Error bars are the standard deviation of at least 10 images. The solid line is a linear fit of the data. **c**, Upper PEG-rich phase prepared at total composition: DEX 6 wt.% and PEG 2.6wt.% + $\lambda$-DNA (48,502 bp) 29 ng/$\mu$L). Scale bar = 10 μm.

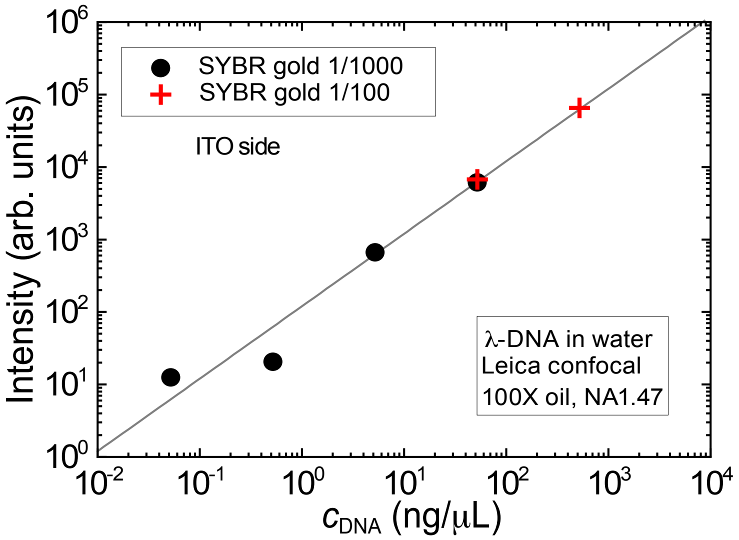


**Fig. S11: DNA concentration dependence of fluorescence intensity from SYBR gold.** The fluorescence intensity of DNA suspension in pure water was measured in confocal microscopy at two conditions, 1/1000 (black circles), and 1/100 (red crosses) of SYBR gold concentrate. Experiments were performed at room temperature (297 ± 0.5 K).

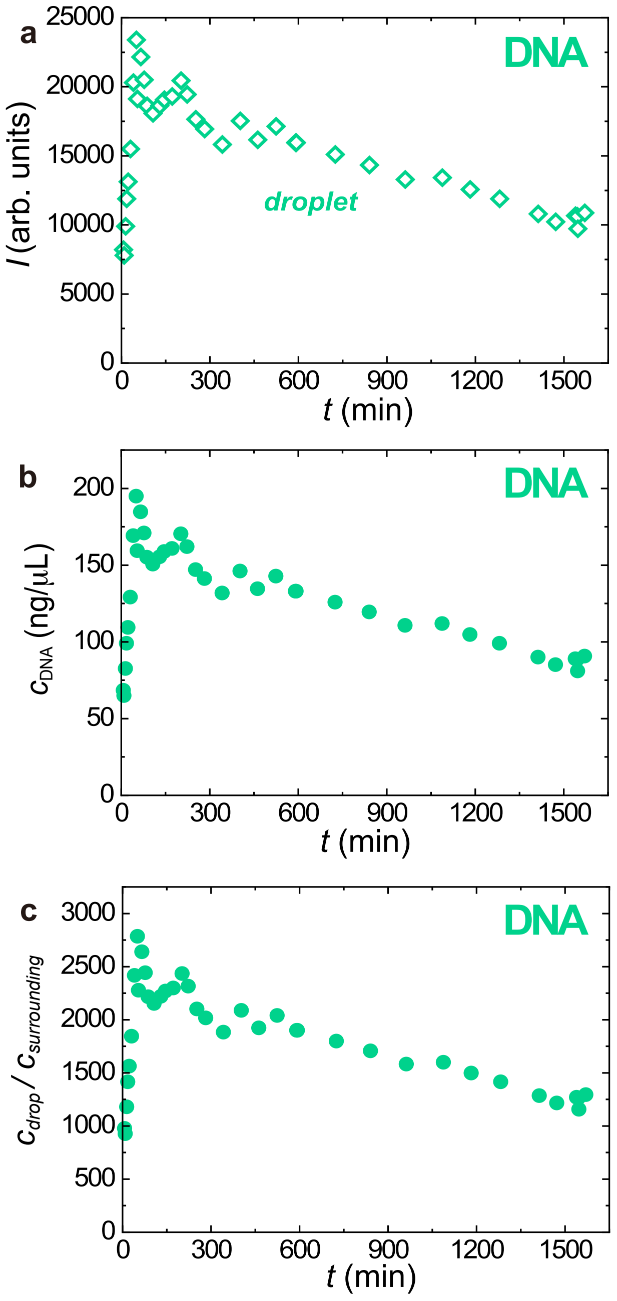


**Fig. S12: Evaluation process of the DNA enrichment factor from confocal data.**

**a,** raw data. **b,** absolute concentration. **c,** concentration ratio.

### 2. DEX concentration ratio

The DEX concentration ratio was estimated as the intensity ratio of the average intensity inside droplet $I_{drop}$ to that of the surrounding phase $I_{surrounding}$, i.e. $I_{drop}/I_{surrounding}$. Before the estimation of the intensity ratio, the background signal in confocal measurements including the dark current of the camera, stray light, and fluorescent signal from substances excluding the target must be subtracted from raw data. The background intensity was measured as the average intensity of pure water in the same sample cell. In time-lapse measurements, a confocal image was obtained by averaging an appropriate minimum number of images for an adequate S/N of 1–16 frames. The average intensity of an image for the emission range to detect DEX is illustrated in Fig. S13a as a function of the number of images for averaging. The intensity is independent of the number of images for averaging.

To calculate $I_{surrounding}$, small ROIs (rectangular regions of approximately 30 μm on each side) were selected from several different positions (approximately 10) from an image, and subsequently, the average intensity and its standard deviation for each ROI were calculated. $I_{surrounding}$ was obtained as the average intensity of those ROIs. The raw data of $I_{drop}$ and $I_{surrounding}$ in a time-lapse measurement are illustrated in Fig. S13b. The background intensity was subtracted from the raw data before calculating the intensity ratio. Finally, the DEX concentration ratio is calculated as shown in Fig. S13c. The experimental error of the intensity ratio was determined by the possible error due to the focal position in the confocal measurements as the factor ± 0.6.

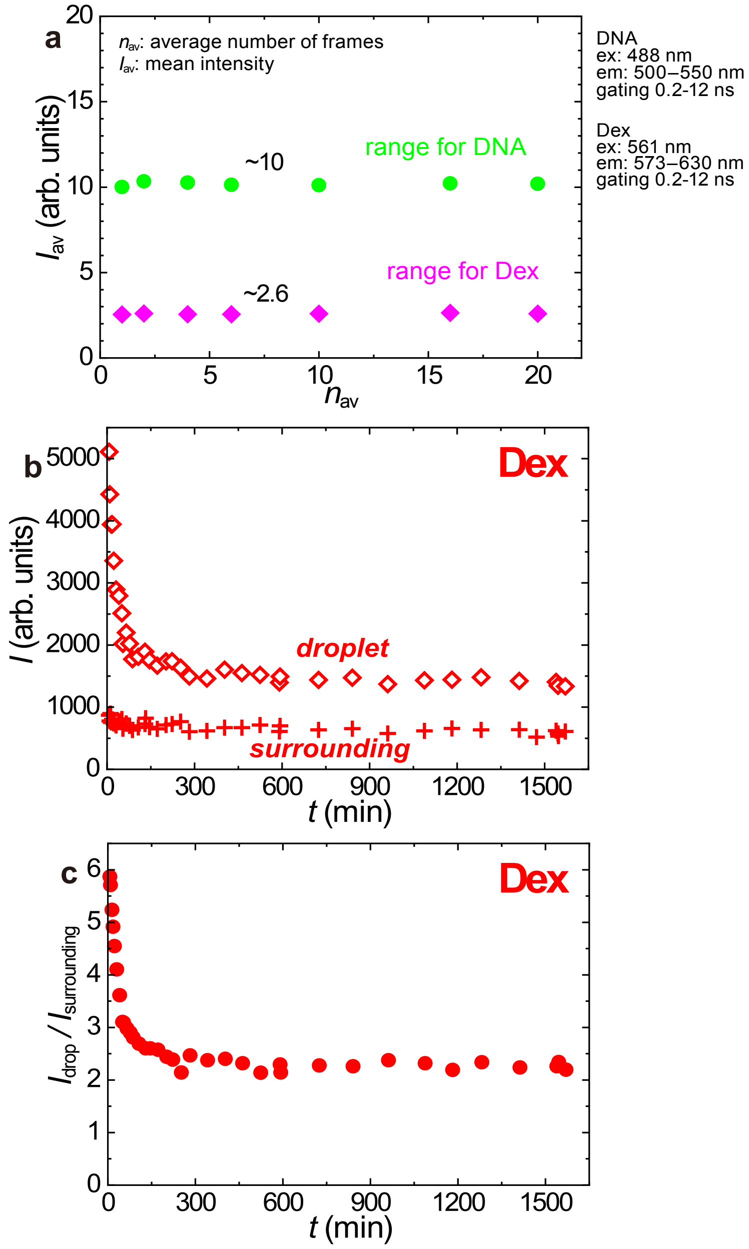


**Fig. S13: Evaluation process of the DEX concentration ratio from confocal data. a,** background signal. **b,** raw data. **c,** intensity ratio.

**Supplementary Text 5: Validity of the temporal change of the intensity ratio**

We checked the possibilities affecting the temporal dependence to confirm that a decrease in the intensity ratio arises from concentration change.

The first possibility is the intensity decrease caused by photobleaching. The contribution of the photobleaching effect to the fluorescence intensity was estimated by the continuous repeat acquisition of spontaneous droplets. These droplets were produced solely by agitating the system in the two-phase region, and therefore, in principle, the DEX concentration inside the droplet should be constant with time. Figure S14a illustrates the average intensity of 139 droplets by repeating the image acquisition 100 times. One round of acquisition took approximately 10 s. The intensity was normalized by the intensity of the first image, and we determined that photobleaching decreased intensity by less than 5 %. In the confocal measurement of the laser-induced droplet, an excitation laser irradiated the sample for 1–20 s for each image. A maximum of 50 timelapse images were obtained. The possible intensity decrease caused by photobleaching in the timelapse measurement was estimated at less than 5 %. The possible photobleaching effect for the DNA is also estimated to be less than 5 % from Fig. S14b.

Next, the signal leakage in multi-wavelength measurement was estimated. Figure S15a illustrates the confocal image of DEX-rich droplets observed in the emission range used for detecting DEX (left) and DNA (right), where the intensity in the right-hand side figure is enhanced by a factor of 25. This sample contained a fluorescent dye (TRITC-DEX 0.2 wt.%) only for DEX without DNA (DEX 6 wt.% (including TRITC-DEX 0.2 wt.%) PEG 2.6wt.%). The right-hand side image highlights the signal leakage to the emission range of the DNA. The relationship between the fluorescence intensity in the emission range of DEX and DNA for the same droplet is presented in Fig. S15a (bottom). The estimated signal leakage from the fluorescent DEX was approximately 4%.

Figure S15b illustrates the results of estimating the signal leakage of SYBR gold (1/1000 of SYBR gold concentrate) to the emission range of DEX, where we observe DEX-rich droplets that do not contain a fluorescent dye for DEX but only for DNA (SYBR gold) (DEX 6 wt.% (TRITC-DEX 0 wt.%) PEG 2.6 wt.% + $\lambda$-DNA (48,502 bp, 29 ng/μL) + 1/1000 of SYBR gold concentrate). The intensity in the left-hand side figure is enhanced by a factor of 1,000. In this case, we observed an intensity distribution; however, the signal leakage was proportional to the original signal. The leakage of the SYBR gold signal was estimated at 0.1%.

Figure S15c and d are raw data of the average intensity of an induced droplet in the timelapse experiment illustrated in Fig. 4 in the main text plotted with the signal leakage of other fluorescent dye estimated from real data. We concluded that the signal leakage was insufficient to cause time dependence.

Finally, we checked if the decrease in the DEX intensity ratio could be attributed to the inappropriate focal position in confocal microscopy. Figure S16 is the side-view (the XZ-slice image) of an induced droplet. The time interval between the two images was 10 min. The diameter is sufficiently large compared to a pixel size, and therefore, the intensity change can be considered a concentration change. There is a clear decrease in the average intensity; i.e. it decreased by a factor of 0.8. From above, we concluded that the DEX concentration ratio of the induced droplets decreased with time.

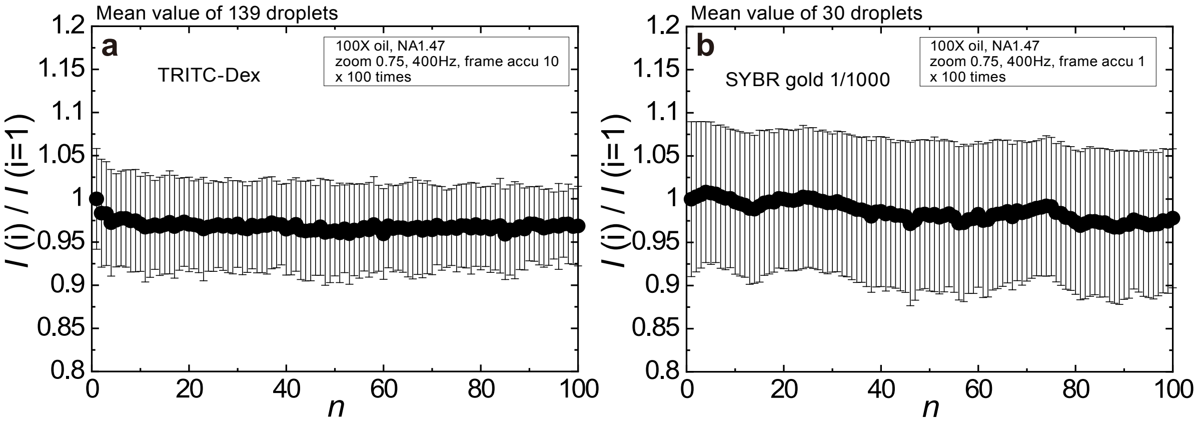


**Fig. S14: Estimation of photobleaching. a,** TRITC-DEX. **b,** SYBR gold.

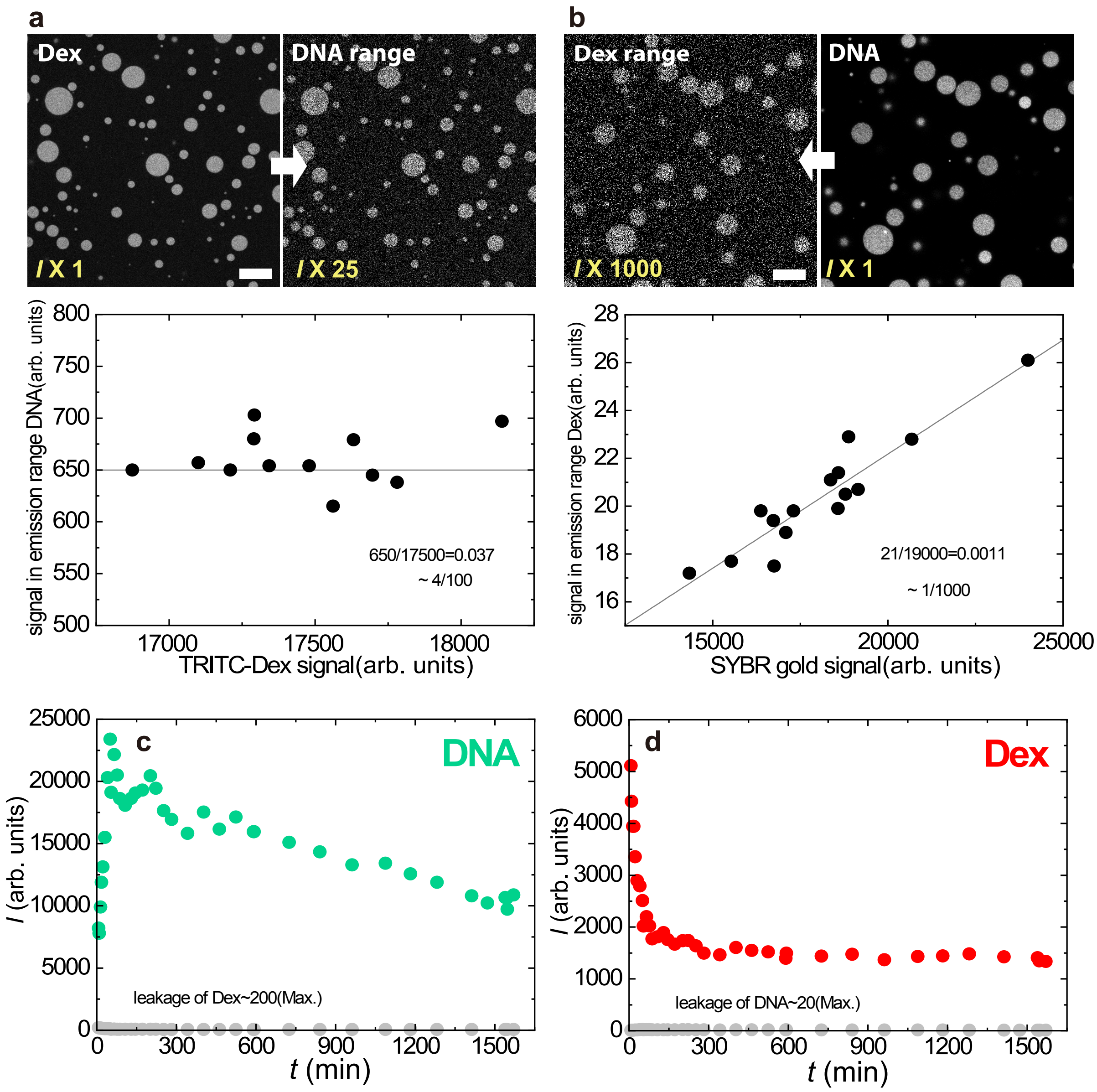


**Fig. S15: Estimation of signal leakage in multi-wavelength detection. a,** Confocal image of droplets visualized by TRITC-DEX (no SYBR gold). Scale bar = 20 μm. **b,** Droplets visualized by SYBR gold (no TRITC-DEX). Scale bar = 20 μm. **c,** Raw fluorescence intensity from DNA plotted with the estimated signal leakage from the DEX signal. d. Raw fluorescence intensity from DEX plotted with the estimated signal leakage from the DNA signal.

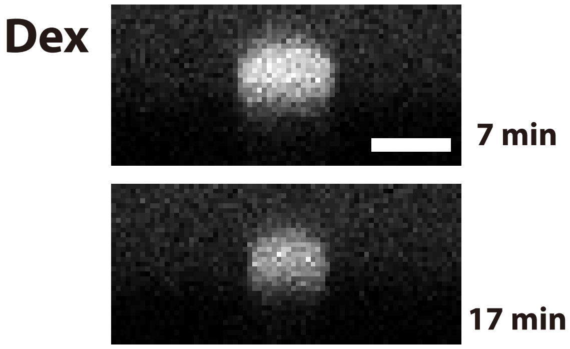


**Fig. S16 : Side view of an induced droplet.** The XZ-slice images of a droplet visualized by fluorescent DEX (TRITC-DEX). The droplet was induced by laser irradiation for 12 min to the upper PEG-rich phase of DEX 6 wt.% PEG 2.6wt.% + $\lambda$-DNA (48,502 bp, 29 ng/μL). (top) $t$ = 7 min. (bottom) $t$ = 17 min after switching off the laser. The intensity ratio between the droplet and the surrounding phase decreased from 7.5 ($t$ = 7 min) to 6 ($t$ = 17 min). Scale bar = 5 μm.

**Supplementary Text 6: DNA enrichment factor and DEX concentration ratio of droplets by spontaneous phase separation**

The DNA enrichment factor and DEX concentration ratio of droplets produced by spontaneous phase separation were estimated under the same condition of confocal microscopy as the laser-induced droplet generation experiments. We prepared a sample of DEX 6 wt.% and PEG 2.6wt.% + $\lambda$-DNA (48,502 bps, 29 ng/μL). Micrometer-size phase-separated droplets were formed by agitating the sample in a conventional vortex mixer. We awaited the sedimentation of droplets for approximately 30 min and extracted the upper PEG-rich phase containing an appropriate number of residual droplets to observe the isolated droplets under a microscope's field of view. For an appropriate comparison with the value obtained in the droplet induction experiment, we placed the sample in the same sample cell used for the droplet generation experiment and awaited the sedimentation of droplets to the ITO surface. Then, we observed the droplets contacted with the ITO surface under the same experimental condition. Typical images of the droplets in Fig. S17. The DNA enrichment factor and DEX concentration ratio were estimated by the same method as in Supplementary Text 3. The results are illustrated in Fig. 5 in the main text.

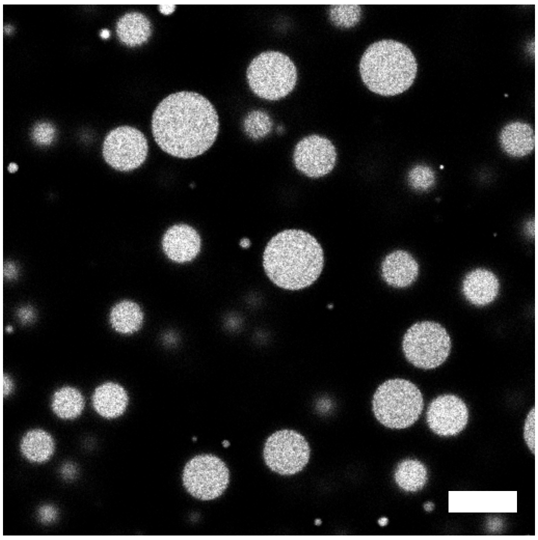

**Fig. S17: Phase-separated DEX-rich droplets obtained by agitation.** Droplets containing DNA were visualized using SYBR gold. DEX 6 wt.% and PEG 2.6wt.% + $\lambda$-DNA (48,502 bp, 29 ng/μL) at 1/1,000 of SYBR gold concentrate. Scale bar = 20 μm.

**Supplementary Text 7: Localization of DNA in a PEG solution**

Duhr and Braun's(3) investigation into DNA suspension under a temperature gradient revealed that the DNA itself exhibits the Soret effect. They reported that the Soret coefficient $S_T$ of the DNA changed its sign around 4 °C and $S_T > 0$ at room temperature (DNA moves away from the hot place). Maeda et al. revealed that, when DNA is added to a PEG solution, its behavior depends on the length of the DNA and PEG concentrations(4, 5). At high concentrations of PEG, the DNA molecules can be localized to the hot region.

To compare the DNA enrichment in the induced phase-separated droplet with the localization of DNA caused by the Soret effect itself(2, 4, 5), we performed local heating experiments in a DNA suspension in a PEG solution without DEX (PEG 3wt.% + DNA), which is approximately the same concentration as that of the system used in the droplet generation experiments (PEG 3.1 wt.%). The temporal evolution of fluorescent images that visualize DNA after laser irradiation for 5 min is illustrated in Fig. S18a. This system did not contain DEX, and it adhered to the condition that phase separation does not occur. Even without phase separation, the DNA molecules were localized at the irradiation spot by local heating at t = 0 s after switching off the laser. The intensity ratio in Fig. S18b was calculated by epifluorescence microscopy (not confocal); this ratio does not correspond to the enrichment factor. However, we can discuss the time scale of diffusion. Note that we performed similar experiments on both DNA in pure water and DNA in the DEX solution at the same concentration as that of each component of the droplet generation experiments. In both cases, the DNA molecules moved away from the hot region created by laser irradiation (Fig. S19).

Further, we performed laser irradiation experiments on a sample of DNA suspension including DEX (DEX 1 wt% PEG 2.4 wt%) that is very close to the composition of the upper PEG-rich phase that is used in the LIPS experiments (DEX 0.7 wt%, PEG 3.1wt%). In that case, DNA formed a ring-shaped localization. But the localisation disappeared in a few minutes (Fig. S20). The concentration increase of DEX due to the Soret effect can be seen, and the concentration distribution disappeared in a few min.

The localization of DNA in the PEG solution is consistent with the observations in the literature(4, 5). However, the temporal evolution was entirely different from the enrichment in a phase-separated droplet. The diffusion of DNA was considerably fast. The intensity homogenized within minutes, which corresponded to a vanishing DNA localization. These results suggest that the long-time localization of DNA in the LIPS droplet induced by the Soret effect is not caused by the high concentration of PEG, and is instead caused by the generation of a phase-separated droplet.

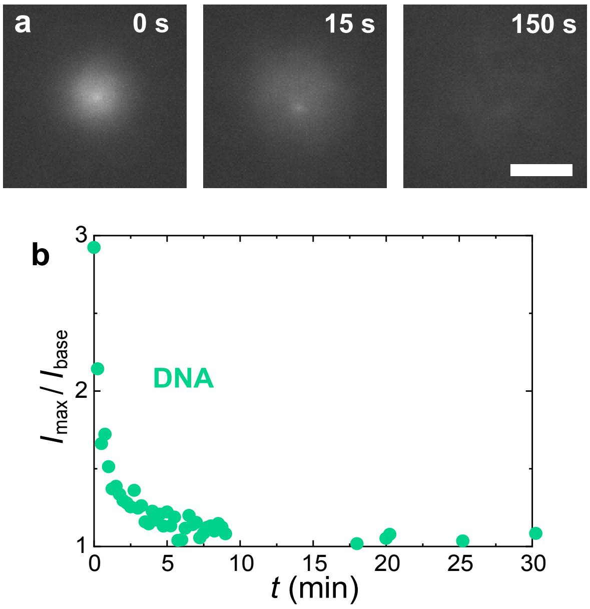


**Fig. S18: Case of local heating in a DNA suspension in a PEG solution (3 wt.%). a,** The temporal evolution in the fluorescent images of the DNA after the laser is disabled. PEG 3 wt.% + $\lambda$-DNA (48,502 bp, 1ng/μL). In this experiment, we used $\lambda$-DNA tagged with a fluorescent dye MFP-488 obtained with nucleic acid labelling reagents (Mirus Bio LLC, label IT, MFP-488). The laser irradiation time was 5 min. The time after the laser was switched off is indicated on the upper right-hand side of each image. DNA is localized at the laser-irradiated position; however, it diffuses rapidly in a few minutes when the laser is disabled. Scale bar = 5 μm. **b,** Temporal evolution of maximum fluorescence intensity normalized by the intensity of the surrounding region. A small bright spot in Fig. S18a is an aggregate or dust, which was excluded from the analysis. The time scale is considerably faster than in the case of DNA enrichment in the phase-separated droplet.

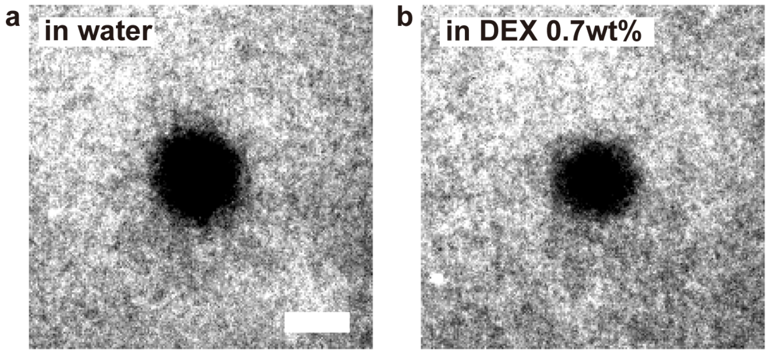


**Fig. S19: Case of local heating in DNA suspensions.** Fluorescent images of DNA visualized by SYBR gold were taken after the laser irradiation for 1 min. Scale bar = 50 μm. **a,** DNA suspension in water. $\lambda$-DNA (48,502 bp, 1.7ng/μL). **b,** DNA suspension in DEX 0.7 wt% solution. $\lambda$-DNA (48,502 bp, 1.7ng/μL).

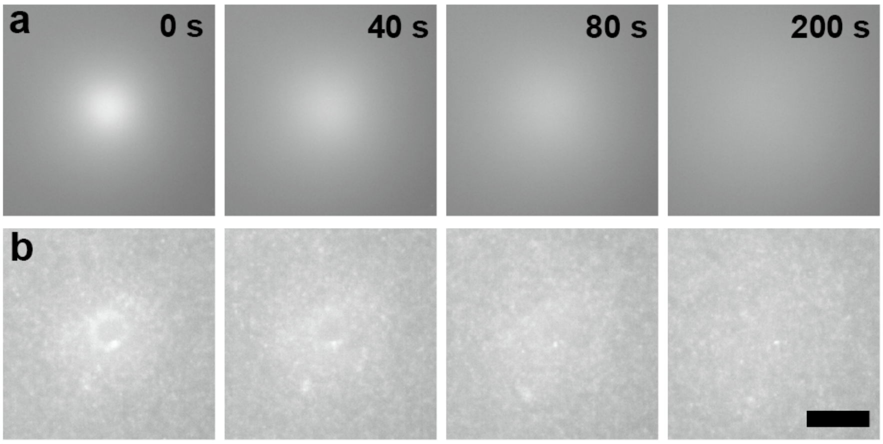


**Fig. S20: Case of a sample in a single-phase region.** Time evolution of the fluorescent image of a sample in one phase region (DEX 1 wt% PEG 2.4 wt% + $\lambda$-DNA (48,502 bp, 1.7ng/μL)) after laser irradiation for 5 min. **a**, DEX **b**, DNA at t = 0 s, 40 s, 80 s, 200 s from the left to the right.
